# Fibroblast activation during decidualization: Embryo-derived TNFα induction of PGI2-PPARδ-ACTIVIN A pathway through luminal epithelium

**DOI:** 10.1101/2022.09.22.509003

**Authors:** Si-Ting Chen, Wen-Wen Shi, Yu-Qian Lin, Zhen-Shang Yang, Ying Wang, Meng-Yuan Li, Yue Li, Ai-Xia Liu, Yali Hu, Zeng-Ming Yang

**Affiliations:** College of Animal Science, Guizhou University, Guiyang 550025, China; College of Veterinary Medicine, South China Agricultural University, Guangzhou, 510642, China; Department of Reproductive Endocrinology, Women’s Hospital, Zhejiang University School of Medicine, 1 Xueshi Road, Hangzhou, Zhejiang, 310006, PR China; Department of Obstetrics and Gynecology, The Affiliated Drum Tower Hospital of Nanjing University Medical School, 321 Zhongshan Rd., Nanjing, 210008, China

**Author notes:** Correspondence: Yali Hu, MD, Department of Obstetrics and Gynecology, Affiliated Drum Tower Hospital, Medical School of Nanjing University, Nanjing 210008, China., Zeng-Ming Yang, Ph.D., College of Animal Science, Guizhou University, Guiyang 550025, China.

**Keywords:** Fibroblast activation, Arachidonic acid, ACTIVIN A, Decidualization

## Abstract

**Objectives:** Human endometrium undergoes cyclical shedding and bleeding, scar-free repair and regeneration in subsequent cycles. Fibroblast activation has been shown to play a key role during normal tissue repair and scar formation. Abnormal fibroblast activation leads to fibrosis. Fibrosis is the main cause of intrauterine adhesion, uterine scaring, and thin endometrium. Endometrial decidualization is a critical step during early pregnancy. There are 75% of pregnancy failures pointed to decidualization defects. Because fibroblast activation and decidualization share similar markers, we assumed that fibroblast activation should be involved in decidualization.

**Materials and Methods:** Both pregnant and pseudopregnant ICR mice were used in this study. Immunofluorescence and immunohistochemistry were applied to examine fibroblast activation-related markers in mouse uteri. Western blotting was used to identify the impact on decidualization. Western blot and RT were used to show how arachidonic acid and its downstream product prostaglandin activate fibroblasts. Additionally, embryo-derived TNFα was shown to stimulate the secretion of arachidonic acid by immunofluorescence, western blot, and ELASA. The aborted decidual tissues with fetal trisomy 16 were compared with control tissues. GraphPad Prism5.0 Student’s t test was used to compare differences between control and treatment groups

**Results:** Fibroblast activation-related markers are obviously detected in pregnant decidua and under in vitro decidualization. ACTIVIN A secreted under fibroblast activation promotes in vitro decidualization. We showed that arachidonic acid released from uterine luminal epithelium can induce fibroblast activation and decidualization through PGI_2_ and its nuclear receptor PPAR-δ. Based on the significant difference of fibroblast activation-related markers between pregnant and pseudopregnant mice, we found that embryo-derived TNFα promotes cPLA_2α_ phosphorylation and arachidonic acid release from luminal epithelium. Fibroblast activation is also detected under human in vitro decidualization. Similar arachidonic acid-PGI_2_-PPARδ-ACTIVIN A pathway is conserved in human endometrium. Compared to controls, fibroblast activation is obviously compromised in human decidual tissues with fetal trisomy 16.

**Conclusions:** Embryo-derived TNFα promotes cPLA2α phosphorylation and arachidonic acid release from luminal epithelium to induce fibroblast activation and decidualization.

**Graphic abstract:** 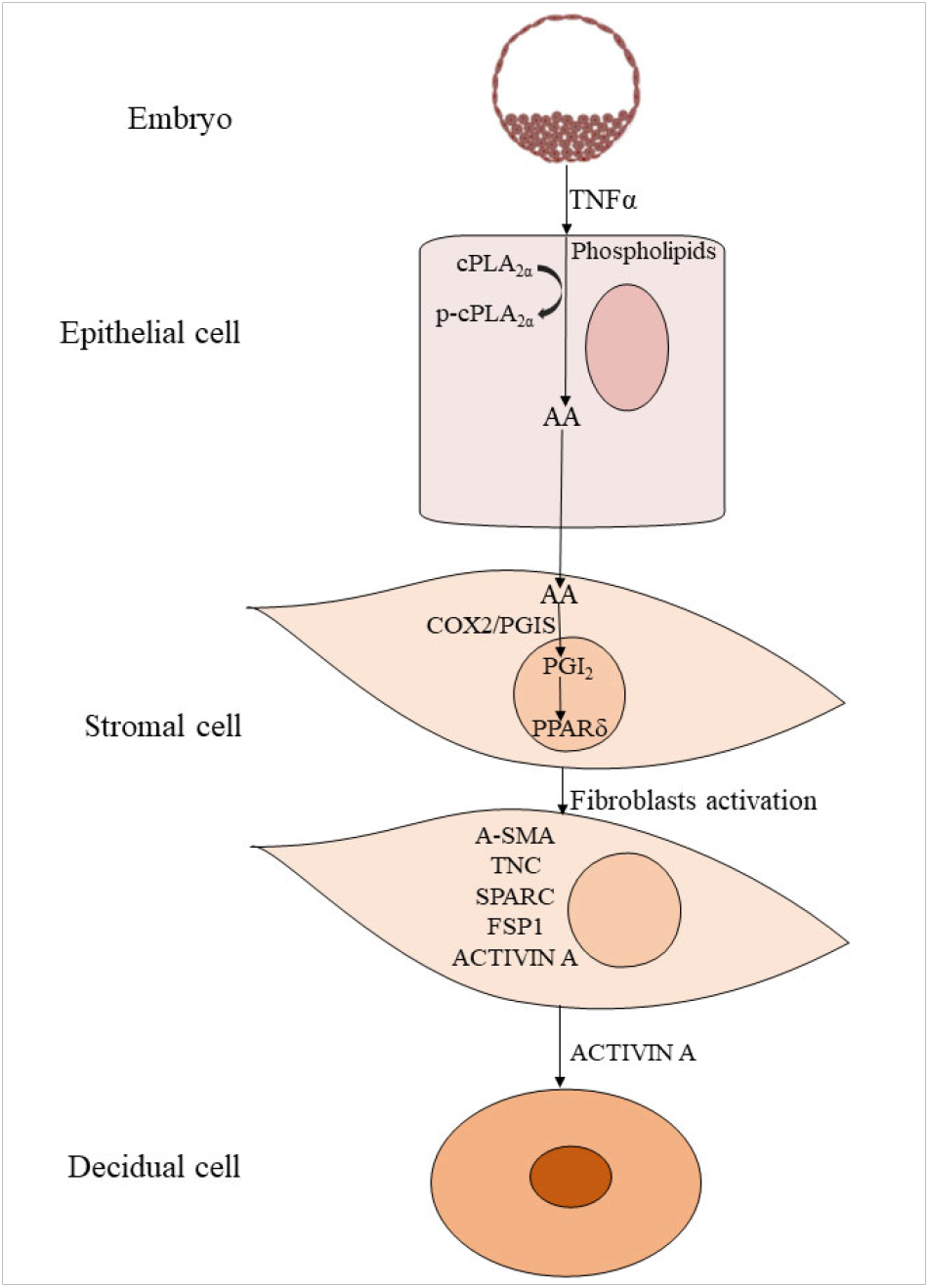

## 1. INTRODUCTION

Fibroblasts are the most numerous cells in connective tissue. Fibroblast activation refers to the process in which dormant quiescent fibroblasts in tissues are stimulated to form functionally related fibroblasts and perform different functions. Fibroblasts synthesize the extracellular matrix (ECM) of connective tissue and play a key role in maintaining the structural integrity of most tissues [1–3]. In healthy and intact tissues, fibroblasts remain a dormant and non-proliferating state. Upon stimulation, dormant fibroblasts acquire contractile properties by inducing the formation of stress fibers, resulting in the formation of myofibroblasts [4; 5]. The myofibroblast is a specialized fibroblast expressing α-smooth muscle actin (α-SMA)[6]. α-SMA is also a marker of cancer-associated fibroblasts (CAFs) [7]. In tumor tissues, activated stromal fibroblasts are called as cancer-associated fibroblasts (CAFs) and show similar characteristics with myofibroblasts [8]. Under the stimulation of cytokines and growth factors, fibroblasts will transform into myofibroblasts at the initial stage and further differentiate to into functionally fibroblasts. Activated fibroblasts are characterized by expressing α-SMA, FAP, Vimentin, Desmin, FSP1, Tenascin C(TNC), periostin (POSTN), and SPARC [9].

In adult tissues, the human endometrium undergoes cyclical shedding and bleeding, scar-free repair and regeneration in subsequent cycles[10; 11]. Activation of fibroblasts plays a key role during normal tissue repair and scar formation [12]. In the uterus, the mesenchyme accounts for about 30-35%, the luminal epithelium and glandular epithelium about 5-10%, and the myometrium about 60-65% of the main uterine cell types. Six cell types have been identified in human endometrium, including stromal fibroblasts, endothelial cells, macrophages, uNK, lymphocytes, epithelial cells, and smooth muscle cells [13]. Based on a recent single-cell transcriptomic analysis of human endometrium, stromal cells were the most abundant cell type in the endometrium [14]. The fertilized egg divides in the fallopian tube to form morula, forming early blastocyst on the fourth day of gestation and entering the uterus [15]. As the blastocyst begins to adhere onto the uterus on day 4 of pregnancy, the activated embryo secretes a series of factors to remodel the stationary fibroblasts through epithelial cells [16]. Endometrial stromal fibroblasts undergo decidualization to form specialized secretory decidual cells, followed by the increase of uterine permeability and immune factors [17]. Decidualization is characterized by significant proliferation, differentiation, and endoreduplication (polyploidy) of endometrial stromal cells near the site of embryo implantation [18]. In mice, decidualization occurs only after the onset of implantation. However, decidualization can also be induced by artificial stimuli [19]. In contrast to mice, human decidualization does not require embryo implantation and is driven by postovulatory progesterone and cAMP [17].

During primate decidualization, αSMA increases at the initial stage and decreases at the final differentiation stage [20]. αSMA is also strongly expressed in rat decidual cells [21]. αSMA expression in interstitial fibroblasts during pregnancy correlates with the onset of the decidual process [22]. Meanwhile, αSMA is strongly expressed in myofibroblast and often recognized as the marker of fibroblast activation [6; 23]. ATP, uric acid and HMGb1 belong to damage associated molecular patterns (DAMPs), which are released following tissue injury or cell death [24]. During the inflammatory phase, the release of these DAMP molecules is able to stimulate the conversion of fibroblasts to αSMA-expressing myofibroblasts [25]. We recently demonstrated that secreted ATP or uric acid under artificial decidualization can induce mouse decidualization [26; 27]. It has been shown that ATP or uric acid can also stimulate fibroblast activation [28; 29]. These studies strongly suggest that there is a similarity between fibroblast activation and decidualization, and fibroblast activation may play an important role during decidualization. Fibrosis is caused by excessive fibroblast activation and the buildup of extracellular matrix and matrix metalloproteinase elements in specific tissues [30]. Endometrial fibrosis is the most common cause of uterine infertility, including implantation failure, and miscarriage [31; 32]. However, whether there is fibrosis during embryo implantation and decidualization and its underlying mechanism are still unclear.

Arachidonic acid (AA) is the biosynthetic precursor of prostaglandins in the cell membrane. cPLA2α is encoded by the *Pla2g4a* gene and a major provider of arachidonic acid. *Pla2g4a* knockout mice show uneven embryo distribution and reduced litter size, suggesting that maternal uterine cPLA2α is critical for successful embryo implantation [33]. Cyclooxygenase COX is the rate-limiting enzyme for prostaglandin synthesis. COX-1 knockout mice are fertile, with only birth defects [34]. COX-2 knockout on C57BL/6J/129 background mice results in impaired implantation, and decidualization, indicating the important role of COX-2 during embryo implantation and decidualization [34]. PGI2 and PGE2 are the two most abundant prostaglandins at the embryo implantation site [35]. PGI2 derived from COX-2 can regulate embryo implantation through PPAR-δ [35]. Both embryonic implantation and decidualization are abnormal in CD1 PPAR-δ-/- mice [36]. However, whether prostaglandins are involved in fibroblast activation during early pregnancy remains unclear.

In this study, our hypothesis was whether fibroblast activation is involved in mouse and human decidualization. The multiple markers of fibroblast activation were detected during mouse in vivo and in vitro decidualization. Embryos-derived TNFα was able to induce the phosphorylation of cPLA2α in luminal epithelium, which liberated arachidonic acid into uterine stroma to promote fibroblast activation through COX2-PGI_2_-PPARδ pathway for inducing decidualization. The underlying mechanism of fibroblast activation was also conserved during human in vitro decidualization. This study offers novel insights into the function of fibroblasts during embryo implantation and decidualization as well as novel approaches to the study of early normal pregnancy and human illnesses.

## 2. MATERIALS AND METHODS

### 2.1. Animals and treatments

Mature CD1 mice (6 weeks of age) was purchased from Hunan Slaike Jingda Laboratory Animal Co., LTD and maintained in specific pathogen free (SPF) environment and controlled photoperiod (light for 12 h and darkness for 12 h). All animal protocols were approved by the Animal Care and Use Committee of South China Agricultural University. In order to induce pregnancy or pseudopregnancy, female mice aged 8-10 weeks were mated with male mice of reproductive age or vasectomized mice (vaginal plug day for day 1). To confirmed the pregnancy of female mice, embryos were flushed from fallopian tubes or uteri from days 1 to 4. On days 4 midnight, 5 and 5 midnight, implantation sites were identified by intravenous injection of 0.1 ml of 1% Chicago blue dye (Sigma-Aldrich, St. Louis, MO) dissolved in saline.

On day 4 of pregnancy (0800-0900 h), pregnant mice were ovariectomized to induce delayed implantation. From days 5 to 7, progesterone was injected daily (1 mg/0.1 ml sesame oil/mice, Sigma-Aldrich) to maintain delayed implantation. Estradiol-17β (1 μg/ml sesame oil/mouse, MCE) was subcutaneously injected on day 7 to activate embryo implantation. Delayed implantation was confirmed by flushing the blastocyst from the uterine horn. The implantation site of the activated uterus was determined by intravenous injection of Chicago blue dye.

### 2.2. Transfer of TNFα-soaked beads

Affi-Gel Blue Gel Beads (Bio-Rad # 1537302) with blastocyst size were incubated with TNFα (0.1% BSA/PBS (410-MT-010, R&D systems, Minnesota, USA) 37°C for 4 h. After washed with M2 (0.1% BSA) for three times, TNFα-soaked beads (15 beads/horn) were transplanted into the uterine horn of day 4 pseudopregnant mice. Beads incubated in 0.1% BSA/PBS were used as control group. Blue bands with beads were identified by injecting Chicago blue into the tail intravenous injection of Chicago blue to observe the implantation site 3 and 24 h after transplantation, respectively.

### 2.3. Immunofluorescence

Immunofluorescence was performed as described previously [37; 38]. Briefly, frozen sections (10 μm) were fixed in 4% paraformaldehyde (158127, Sigma Aldrich, St. Louis, MO) for 10 min.Frozen or paraffin sections were blocked with 10% horse serum for 1 h at 37 °C and incubated overnight with appropriate dilutions of primary antibodies at 4°C. The primary antibodies used in this study included anti-α-SMA (19245T, Cells Signaling Technology, Danvers,MA), anti-SPARC (8725S, Cells Signaling Technology), anti-FSP1 (13018S, Cells Signaling Technology), anti-POSTN (SAB2101847, Sigma-Aldrich), anti-P-CPLA2α (2831S, Cells Signaling Technology). After washing in PBS, sections were incubated with secondary antibody (Jackson ImmunoResearch, West Grove, PA) for 40 min, counterstained with 4, 6- diamidino-2-phenylindole dihydrochloride (DAPI, D9542, Sigma-Aldrich) or propidium iodide (PI) and were mounted with ProLong™ Diamond Antifade Mountant (Thermo Fisher, Waltham, MA). The pictures were captured by laser scanning confocal microscopy (Leica, Germany).

### 2.4. Immunohistochemistry

Immunohistochemistry was performed as described previously [39]. In short, paraffin sections (5 μm) were deparaffined, rehydrated, and antigen retrieved by boiling in 10 mM citrate buffer for 10 min. Endogenous horseradish peroxidase (HRP) activity was inhibited with 3% H_2_O_2_ solution in methanol. After washing for three times with PBS, the sections were incubated at 37°C for 1 h in 10% horse serum for blocking, incubated overnight in each primary antibody at 4°C. The primary antibodies used in this study included anti-α-SMA, anti-TNC (ab108930, Abcam, Cambridge, UK). anti-FSP1, and anti-POSTN. After washing, the sections were incubated with biotinylated rabbit anti-goat IgG antibody (Zhongshan Golden Bridge, Beijing, China) and streptavidin-HRP complex (Zhongshan Golden Bridge). According to the manufacturer’s protocol, the positive signals were visualized using DAB Horseradish Peroxidase Color Development Kit (Zhongshan Golden Bridge). The nuclei were counter-stained with hematoxylin.

### 2.5. siRNA transfection

The siRNAs for mouse Activin a were designed and synthesized by Ribobio Co., Ltd. (Guangzhou, China). Following manufacturer’s protocol, cells were transfected with each *Inhba* siRNA using Lipofectamine 2000 Transfection Reagent (Invitrogen, Grand Island, NY) for 6 h. The siRNA sequences were listed in Table 1.

**Table 1.**
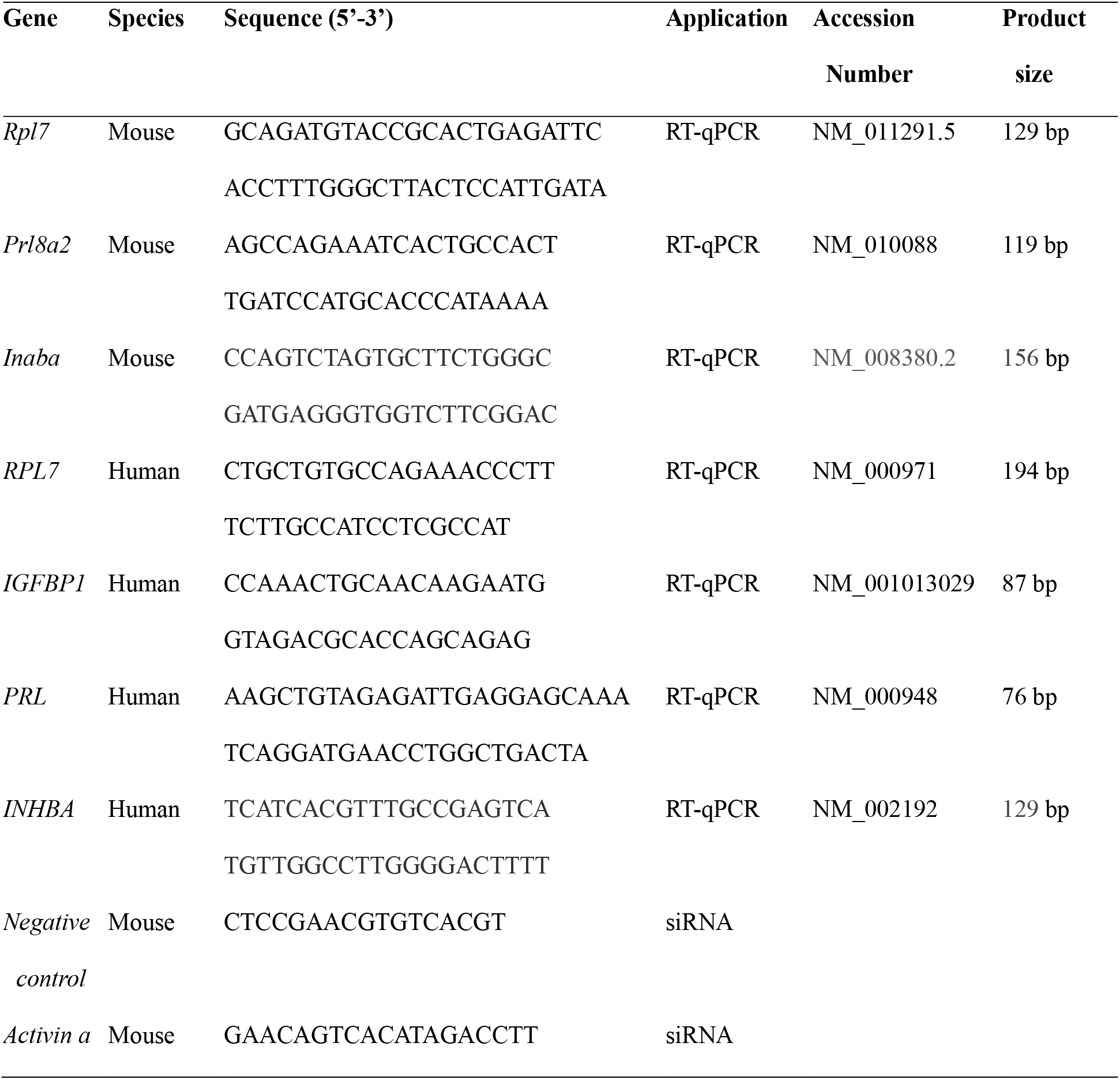
Primers and siRNA sequences used in this study.

### 2.6. Arachidonic acid assay

The arachidonic acid ELISA kit was used to detect arachidonic acid in the supernatant according to the manufacturer’s instructions (Elabscience, E-EL-0051c, Wuhan, China). This kit’s sensitivity is greater than 0.94ng/ml. In brief, 50 μl of each sample was incubated at 37°C for 45 minutes with 50 μl of biotinylated antibody working solution, 100 μL of HRP enzyme conjugate working solution for 30 minutes, and 90 μL of substrate solution for 15 minutes before being stopped with 50 μL of substrate solution. The solution was immediately read at 450 nm with a Biotek microplate reader (ELX808). Absorbance values for arachidonic acid standards were calculated in the same way. The arachidonic acid standard curve was used to calculate the concentrations of arachidonic acid.

### 2.7. Isolation and treatment of mouse uterine luminal epithelial cells

Uterine luminal epithelial cells were isolated as previously described [40]. The uteri from the estrous mice or day 4 of pseudopregnancy were cut longitudinally, washed in HBSS, incubated in 0.2% (W/V) trypsin (0458, Amresco, Cleveland, USA) and 6 mg/ml dispase (Roche Applied Science, 4942078001, Basel, Switzerland) in 4.3 mL HBSS for 1.5 h at 4°C, 30 min at room temperature, and 10 min at 37°C. After rinsing in HBSS, the epithelial cells were precipitated in 5% BSA in HBSS for 7 min. After the collected epithelial cells were cultured in DMEM/F12 (D2906, Sigma-Aldrich) with 10% FBS in a culture plate for 30 min, the unattached epithelial cells were transferred into new culture plates for further culture. Luminal epithelial cells were treated with TNFα (410-MT-010, R&D systems) in DMEM/F12 with 2% charcoal-treated FBS (cFBS, Biological Industries, Cromwell, CT).

### 2.8. Isolation and treatment of mouse endometrial stromal cells

Mouse endometrial stromal cells were isolated as previously described [38]. Briefly, mouse uteri on day 4 of pseudopregnancy were cut longitudinally, washed in HBSS, and incubated with 1% (W/V) trypsin and 6 mg/ml dispase in 3.5 mL HBSS for 1 h at 4 °C, for 1h at room temperature and for 10 min at 37°C. The uterine tissues were washed with Hanks’ balanced salt solution, incubated in 6 ml of HBSS containing 0.15 mg/ml Collagenase I (Invitrogen, 17100-017) at 37°C for 35 min. Primary endometrial stromal cells were cultured with DMEM/F12 containing 10% heat-inactivated fetal bovine serum (FBS).

Mouse in vitro decidualization was performed as previously described [41]. Primary endometrial stromal cells were treated with 10 nM of Estradiol-17 β and 1 μM of P4 in DMEM/F12 containing 2% charcoal-treated FBS (cFBS, Biological Industries) to induce decidualization in vitro for 72 h. Stromal cells were treated with TNC(3358-TC-050, R & D systems), FSP1(HY-P71084, MedChemExpress, NJ, USA), arachidonic acid (A3611,Sigma-Aldrich), PGI analogue ILOPROST (HY-A0096, MedChemExpress), PPAR-δ agonist GW501516 (317318-70-0, Cayman Chemical), COX-2 antagonist NS 398(S8433, Selleck, Shanghai. China), PPAR-δ antagonist GSK0660 (1014691-61-2, Selleck), and ACTIVIN A (HY-P70311, MedChemExpress) in DMEM/F12 containing 2% carbonate-treated FBS, respectively.

### 2.9. Culture and treatment of human cell lines

Ishikawa endometrial adenocarcinoma cells line (Chinese Academy of Science, Shanghai, China) and human endometrial stromal cell 4003 (ATCC, CRL-4003) (American Type Culture Collection) were cultured in DMEM/F12 with 10% FBS, and supplemented with 100 units/ml penicillin and 0.1 mg/ml streptomycin (PB180429, Procell, Wuhan, China) at 37°C, 5% CO2. TNFα was used to treat Ishikawa cells.

### 2.10. Co-culture of epithelial cells and stromal cells

The uterine luminal epithelial cells isolated from mouse uteri on day 4 of pseudopregnancy were cultured to confluence on coverglasses. The endometrial stromal cells isolated from mouse uteri on day 4 pseudopregnancy were cultured in a culture plate with four plastic pillars. Then the coverglasses with epithelial cells were transferred onto the four plastic pillars of culture plates with stromal cells for further culture and treatments. The co-culture of human ISHIKAWA cells with human stromal 4003 cells was performed as in mice.

### 2.11. Western blot

Western blot was performed as previously described [37]. The primary antibodies used in this study included phosphorylated CPLA_2α_ (SC-438, Santa Cruz Biotechnology), CPLA_2α_, TNC, SPARC (8725S, Cells Signaling Technology), α-SMA, COX-2 (12282T, Cells Signaling Technology), PGIS (100023, Cayman Chemical), PPARδ (ab178866, Abcam), BMP2 (A0231, Abclonal, Wuhan, China), WNT4 (sc-376279, Santa Cruz Biotechnology, Dallas, TX), E2F8 (A1135, Abclonal), CYCLIN D3 (2936T, Cells Signaling Technology), TUBULIN (2144S, Cells Signaling Technology), GAPDH (sc-32233, Santa Cruz Biotechnology). After the membranes were incubated with an HRP-conjugated secondary antibody (1:5000, Invitrogen) for 1 h, the signals were detected with an ECL Chemiluminescent Kit (Millipore, USA).

### 2.12. Real-time RT-PCR

The total RNA was isolated using the Trizol Reagent Kit (9109, Takara, Japan), digested with RQ1 deoxyribonuclease I (Promega, Fitchburg, WI), and reverse-transcribed into cDNA with the Prime Script Reverse Transcriptase Reagent Kit (Takara, Japan). For real-time PCR, the cDNA was amplified using a SYBR Premix Ex Taq Kit (TaKaRa) on the CFX96 Touch™ Real-Time System (Bio-Rad). For real-time PCR System (Bio-Rad). Data were analyzed using the 2-ΔΔCt method and normalized to *Rpl7* (mouse) or *RPL7* (human) level. The corresponding primer sequences of each gene were provided in Table 1.

### 2.13. Collection of human decidual tissues with fetal trisomy 16

Endometrial decidual tissues were taken from women between the ages of 31 and 38 who underwent voluntary pregnancy termination (8-10 weeks gestation) at Hangzhou Women’s Hospital and Drum Tower Hospital affiliated with Nanjing University School of Medicine in China. To diagnose aneuploidy, chromosomal microarray (CMA) was used to test the fetal tissues of patients. In this study, decidual tissues were collected after fetal tissues were identified as trisomy 16. The decidual tissues were collected from women who decided to terminate their pregnancy due to an unplanned pregnancy. For further examination, all samples were rinsed with saline to eliminate excess blood, fixed in 10% PBS-buffered formalin, and embedded in paraffin. The informed consents were obtained from all the patients prior to the collection of samples. This study was approved by The Ethics Committee of Zhejiang University School of Medicine’s Obstetrics and Gynecology Hospital and the Human Research Committee of Nanjing Drum Tower Hospital, respectively.

### 2.14. Statistical analysis

The data were analyzed by GraphPad Prism5.0 Student’s T test was used to compare differences between two groups. The comparison among multiple groups was performed by ANOVA test. All the experiments were repeated independently at least three times. In the mouse study, each group had at least three mice. Data were presented as mean ± standard deviation (SD). A value of p<0.05 was considered significant.

## 3. RESULTS

### 3.1. Fibroblast activation is present during mouse early pregnancy

Alpha SMA, SPARC, FSP1, TNC, and SPARC are well recognized markers of fibroblast activation [9; 42]. Immunofluorescence and immunohistochemistry showed that TNC, SPARC, and FSP1 were mainly localized in the primary decidual region after embryo implantation. SPARC immunofluorescence was strongly observed in subluminal stromal cells from day 4 midnight to day 5 midnight of pregnancy. However, TNC immunostaining was strongly detected in subluminal stromal cells on day 4 morning, day 4 midnight and day 5 of pregnancy, but disappeared on D5 midnight. FSP1 immunostaining wasn’t detected in mouse uterus on day 4 morning and day 4 midnight of pregnancy, but detected in subluminal stromal cells at implantation sites on day 5 morning and day 5 midnight of pregnancy (Fig. 1A). Under in vitro decidualization, the protein levels of αSMA, TNC and SPARC were significantly increased compared with the control group (Fig. 1B-source data-1). These results suggested that fibroblast activation should be present under in vivo and in vitro decidualization.

**Fig. 1.**
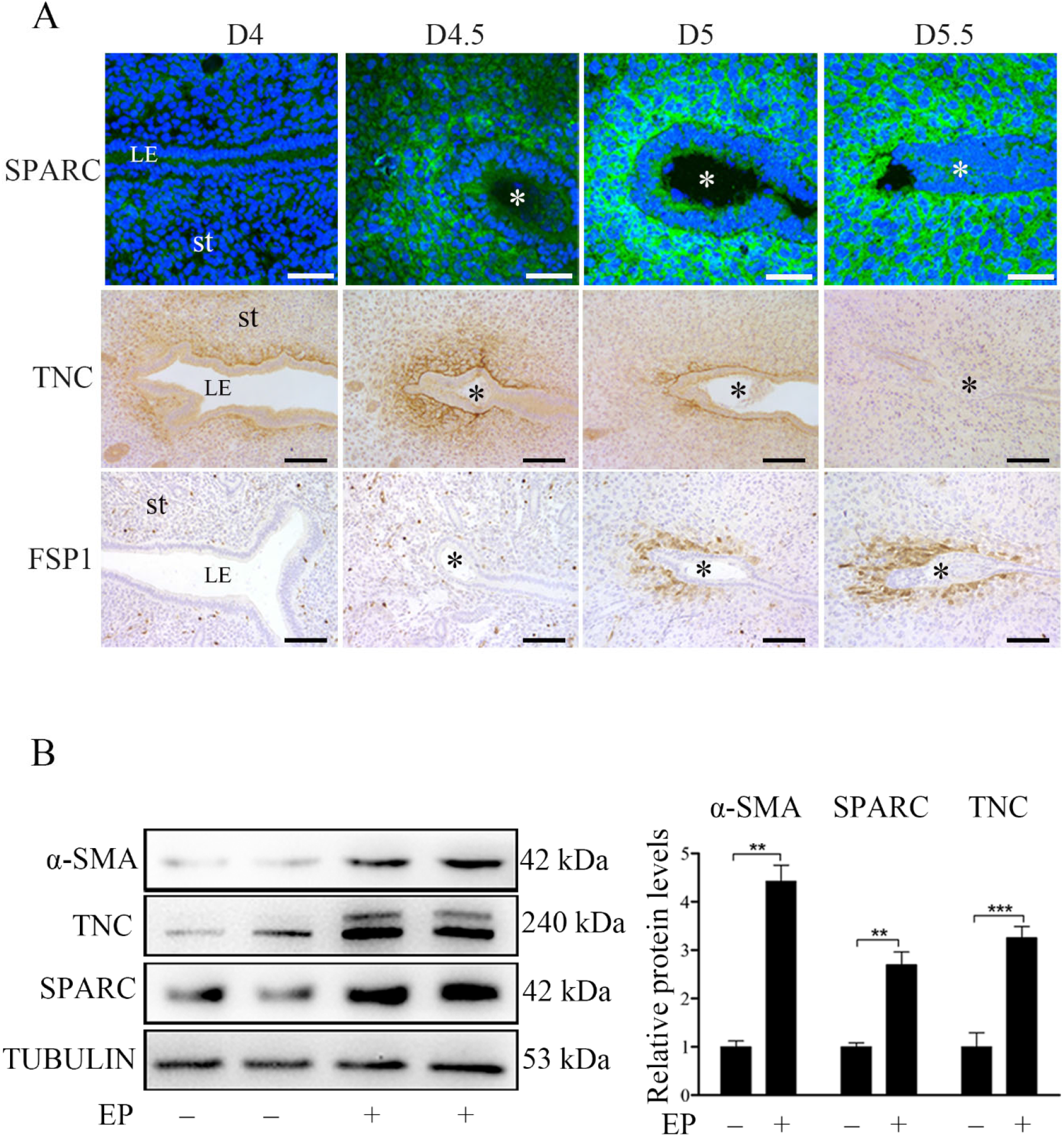
The protein localization and levels of markers of fibroblast activation in mouse uteri during early pregnancy. (**A**) Immunofluorescence of SPARC and POSTN, and immunohistochemistry of TNC and FSP1 in mouse uteri on day 4 0900 (D4, n=5), day 4 24:00 (D4.5, n=5), day 5 0900 (D5, n=5), and day 5 2200 (D5.5, n=5) of pregnancy. Three mice are used in each group. LE, luminal epithelium; St, stroma; * Embryo. Scale bar, 50 μm. (**B**) Western blot analysis of α-SMA, SPARC, TNC protein level under in vitro decidualization (EP) for 24 h. *, p < 0.05; **, p < 0.01; ***, p < 0.001.

### 3.2. Fibroblasts activation promotes decidualization by secreting ACTIVIN A

BMP2 and WNT4 are essential to mouse decidualization [43; 44]. E2F8 and CYCLIN D3 are markers of polyploidy during mouse decidualization [45]. It has been shown that activated fibroblasts can secrete FSP1, SPARC, TNC, and ACTIVIN A [9; 46]. In order to examine whether fibroblast activation is involved in mouse decidualization, mouse stromal cells were treated with FSP1, SPARC, TNC, and ACTIVIN A to induce in vitro decidualization, respectively. TNC treatment upregulated the protein levels of WNT4 and CYCLIN D3, but had no obvious effects on BMP2 and E2F8 (Fig. 2A-source data-1). FSP1 treatment significantly increased BMP2 and WNT4 protein levels, but had no obvious effects on E2F8 and CYCLIN D3 (Fig. 2B-source data-2). SPARC overexpression in mouse stromal cells also upregulated BMP2 and CYCLIN D3 protein levels, but had no significant effect on WNT4 and E2F8 (Fig. 2C-source data-3). Because ACTIVIN A is secreted under fibroblast activation [47], we examined ACTIVIN A protein levels in the uteri on D4 09:00 and 24:00 of pregnancy and pseudopregnant mice, respectively. The protein levels of ACTIVIN A on D4 09:00 and 24:00 of pregnancy were significantly higher than that in pseudopregnant mice. The protein level of ACTIVIN A on day 4 24:00 was significantly higher than that on day 4 09:00 (Fig. 2D-source data-4), indicating that the secretion of ACTIVIN A increased after embryo implantation. After mouse stromal cells were treated with ACTIVIN A, all the protein levels of BMP2, WNT4, E2F8, and CYCLIN D3 were obviously up-regulated (Fig. 2E-source data-5). Under in vitro decidualization, ACTIVIN A treatment also significantly increased all the protein levels of BMP2, WNT4, E2F8, and CYCLIN D3 (Fig. 2F-source data-6). These results suggest a positive correlation between ACTIVIN A and decidualization.

### 3.3. PGI_2_ promotes fibroblast activation and decidualization through PPAR-δ pathway

Although we just showed the presence of fibroblast activation during decidualization, what initiates fibroblast activation is still unknown. A previous study indicated that PGE_2_ and PGI_2_ are the most abundant prostaglandins at implantation sites in mouse uterus [35]. Both PGE_2_ and PGI_2_ are essential to mouse decidualization [35; 48]. When mouse stromal cells were treated with PGE2, PGE2 had no obvious effects on the protein levels of TNC, αSMA and SPARC (Fig. 3A-source data-1). However, ILOPROST, a PGI2 analog, significantly up-regulated the protein levels of TNC, αSMA and SPARC. Peroxisome proliferator-activated receptor δ (PPARδ), the PGI_2_ nuclear receptor, was also significantly increased by ILOPROST (Fig. 3B-source data-2). Further analysis showed that PPAR-δ agonist GW501516 was also able to upregulate the protein levels of TNC, αSMA and SPARC (Fig. 3C-source data-3). After stromal cells were treated with either ILOPROST or GW501516, decidualization markers (BMP2, WNT4, E2F8 and CYCLIN D3) were also significantly up-regulated (Fig. 3D-source data-4, E-source data-5). ILOPROST treatment also obviously upregulated the protein level of ACTIVIN A (Fig. 3F-source data-6). These results showed that PGI2 initiated fibroblast activation and promoted decidualization through the PPAR-δ pathway.

### 3.4. Arachidonic acid induces the fibroblasts activation and promotes decidualization through stimulating Activin A secretion

Arachidonic acid is the precursor of prostaglandin biosynthesis and can be liberated from membrane lipids through the phosphorylation of cPLA2α [49]. cPLA2α is significantly expressed in the luminal epithelium in mouse uterus during peri-implantation period and essential for mouse embryo implantation [33], suggesting that arachidonic acid should be released from luminal epithelium. After stromal cells were treated with arachidonic acid, the markers of fibroblast activation (TNC, αSMA and SPARC) were significantly up-regulated (Fig. 4A-source data-1). Treatment with arachidonic acid also upregulated the protein levels of decidualization markers (BMP2, WNT4, E2F8 and CYCLIN D3) (Fig. 4B-source data-2) and the mRNA level of *Prl8a2* (another marker for mouse decidualization) (Fig. 4D). Under in vitro decidualization, arachidonic acid treatment also significantly increased the decidualization markers (Fig. 4C-source data-3). These results indicated that arachidonic acid might promote decidualization through fibroblast activation. Additionally, treatment with arachidonic acid distinctly stimulated the protein levels of COX-2, PGIS and PPAR-δ, but had no obvious effects on PGES (Fig. 4E-source data-4). The stimulation of arachidonic acid on PGIS and PPARδ was abrogated by COX-2 inhibitor NS398 (Fig. 4F-source data-5). The induction of arachidonic acid on *Prl8a2* mRNA levels was also suppressed by either COX-2 inhibitor NS398 or PPAR-δ antagonist GSK0660 (Fig. 4H, 4I). Arachidonic acid also significantly induced the protein level of ACTIVIN A and the mRNA level of *Inhba* (encoded for ACTIVIN A) (Fig. 4G, 4J-source data-6). Furthermore, the induction of arachidonic acid on *Prl8a2* expression was abrogated by either ACTIVIN A inhibitor (SB431542) or *Inhba* siRNA (Fig. 4K, 4L). These results suggested that arachidonic acid induced fibroblast activation and promoted decidualization through ACTIVIN A.

**Fig. 2.**
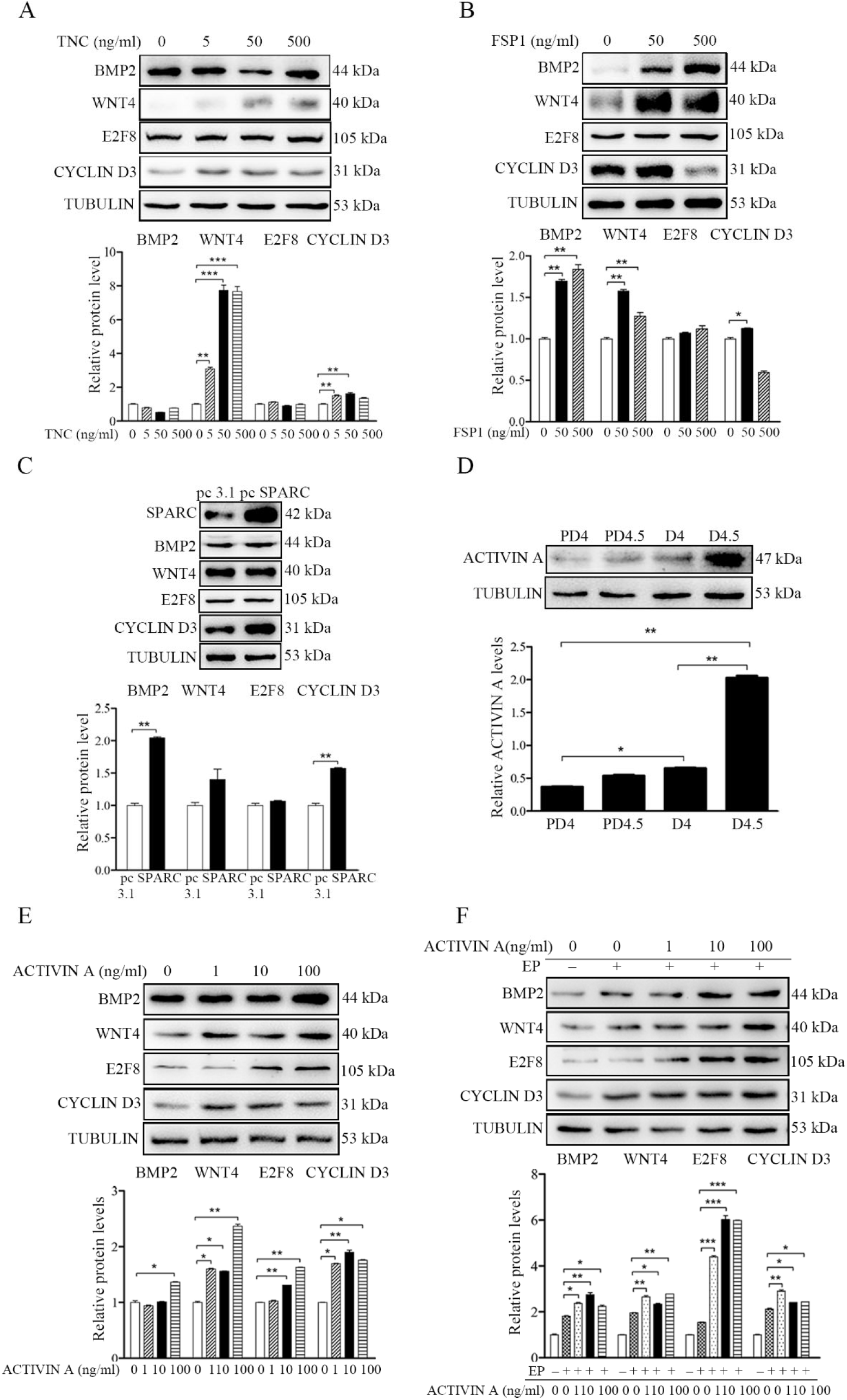
Fibroblasts activation promotes decidualization by secreting ACTIVIN A. (**A**) Western blot analysis on the effects of TNC on decidualization markers (BMP2, WNT4, E2F8 and CYCLIN D3) after stromal cells were treatment with TNC for 72 h. (**B**) Western blot analysis of the effects of FSP1 on decidualization markers after stromal cells were treated with FSP1 for 72 h. (**C**) Western blot analysis on the effects of Sparc overexpression on decidualization markers after overexpression of *Sparc* gene in cultured stromal cells. (**D**) Western blot analysis on ACTIVIN A protein levels in mouse uteri on day 4 0900 and day 4 2400 of pregnancy and pseudopregnancy, respectively. (**E**)Western blot analysis on the effects of ACTIVIN A on decidualization markers after stromal cells were treated with ACTIVIN A for 72 h. (**F**) Western blot analysis on the effects of ACTIVIN A on decidualization markers after stromal cells were treated with ACTIVIN A for 48 h under in vitro decidualization. All data were is presented as means ± SD.

**Fig. 3.**
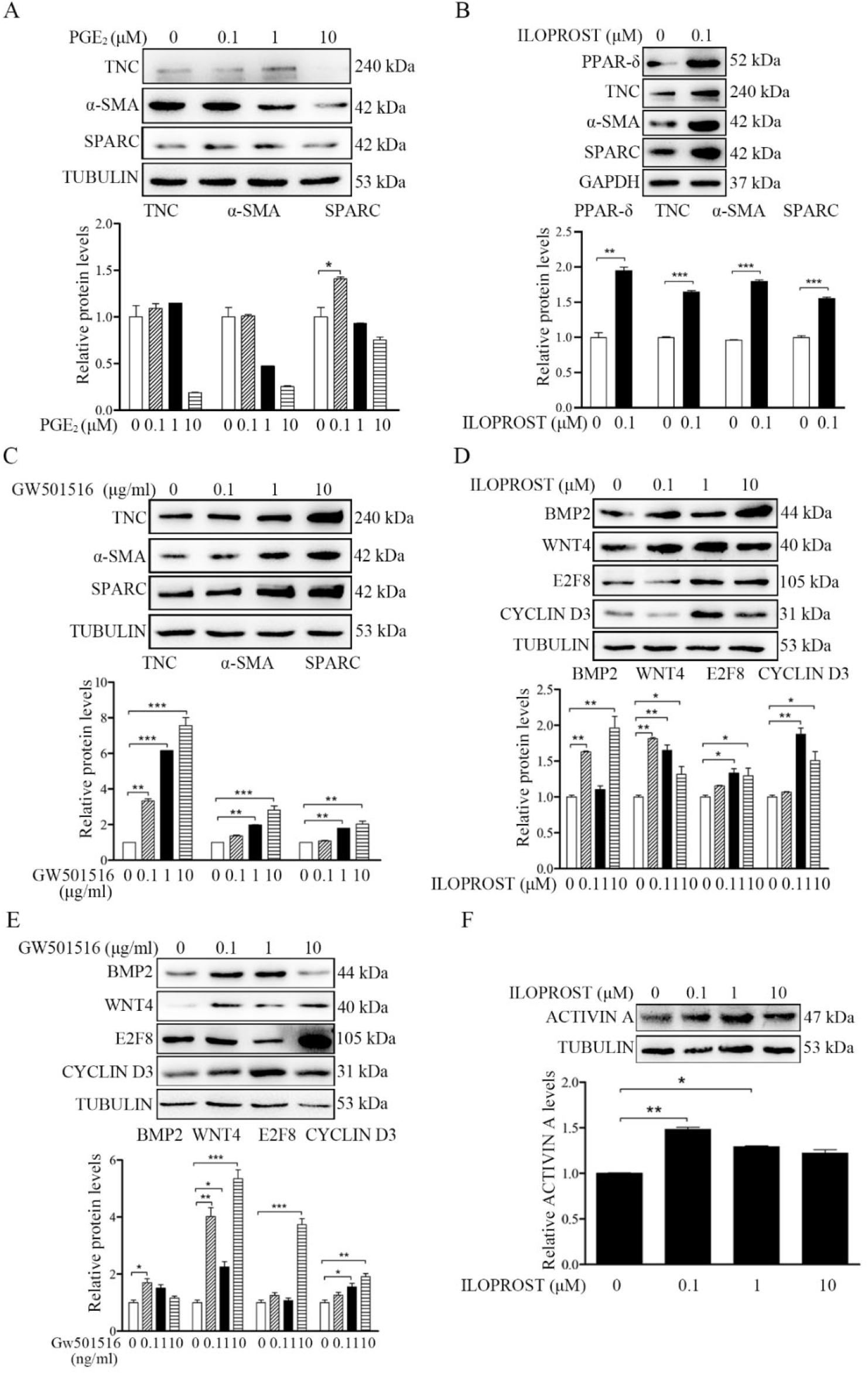
Western blot analysis on effects of prostaglandins on fibroblast activation and decidualization in mouse stromal cells. (**A**) The effects of PGE2 on markers of fibroblast activation. (**B**) The effects of ILOPROST, PGI_2_ analog, on markers of fibroblast activation after stromal cells was treated with PGI_2_ for 12 h. (**C)** The effects of GW501516, PPARδ agonist, on markers of fibroblast activation. (**D**) The effects of ILOPROST on decidualization markers. (**E**) The effects of GW501516 on decidualization markers. (**F**) The effects of ILOPROST on ACTIVIN A protein levels after stromal cells were treated with ILOPROST for 24 h.

**Fig. 4.**
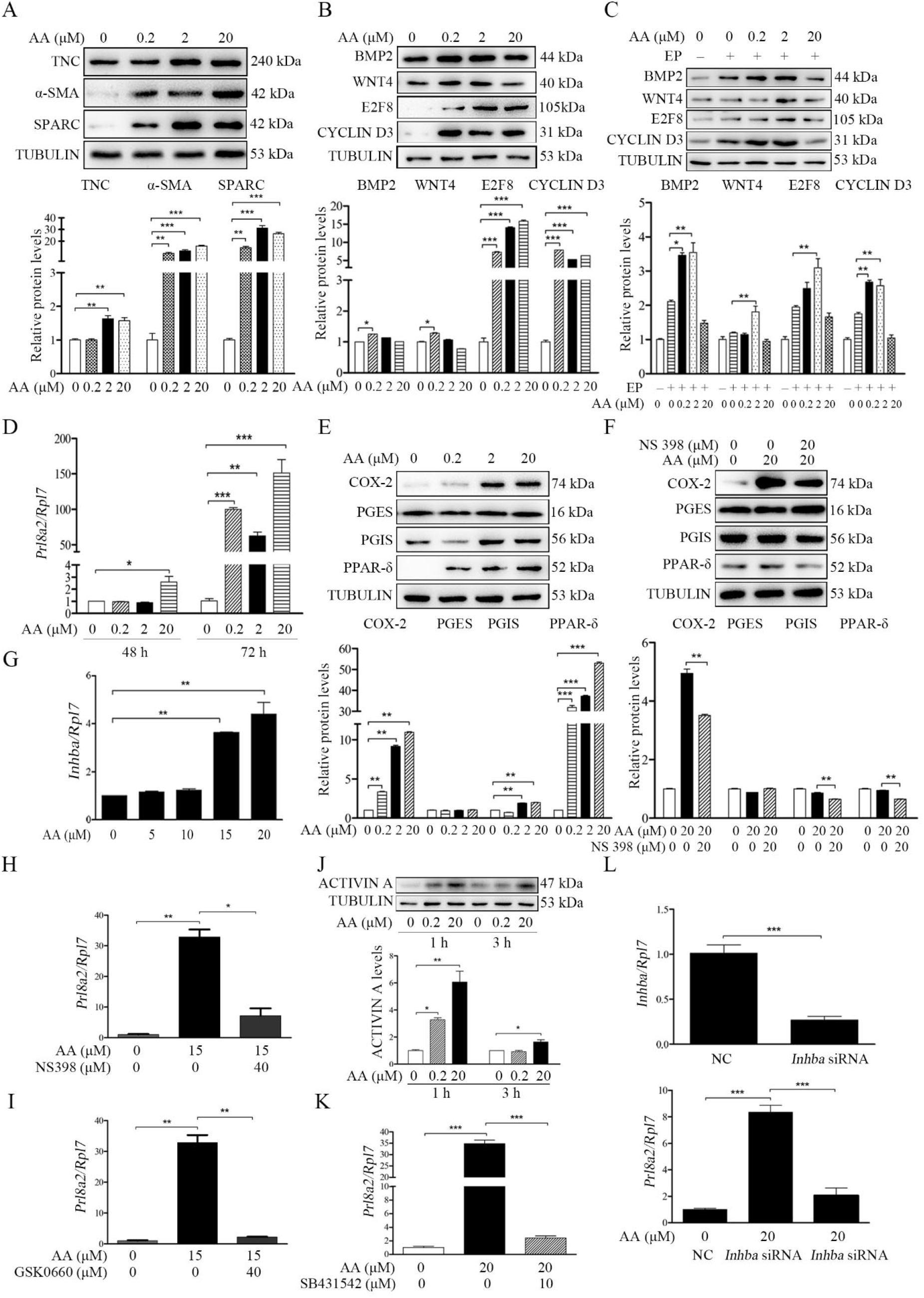
Effects of arachidonic acid on fibroblast activation and decidualization through PGI-PPARδ-ACTIVIN A pathway. (**A**) Western blot analysis on effects of arachidonic acid on markers of fibroblast activation after stromal cells were treated with arachidonic acid for 6 h. (**B**) Western blot analysis on effects of arachidonic acid on decidualization markers after stromal cells were treated with arachidonic acid for 48 h. (**C**) Western blot analysis on effects of arachidonic acid on decidualization markers after stromal cells were treated with arachidonic acid for 48 h under in vitro decidualization. EP, 17β-estradiol + progesterone. (**D**) QPCR analysis of *Prl8a2* mRNA level after stromal cells were treated with arachidonic acid for 72 h. (**E**) Western blot analysis on effects of arachidonic acid on COX2, PGES, PGIS and PPARδ protein levels after stromal cells were treated with arachidonic acid for 6 h. (**F**) Western blot analysis on effects of NS398 (COX-2 inhibitor) on arachidonic acid induction of COX2, PGES, PGIS and PPARδ protein levels after stromal cells were treated with arachidonic acid for 48 h in the absence or presence of NS398. (**G**) QPCR analysis of *Inhba* mRNA level after stromal cells were treated with arachidonic acid for 72 h. (**H**) QPCR analysis of on effects of NS398 on arachidonic acid induction of *Prl8a2* mRNA level after stromal cells were treated with arachidonic acid for 72h. (**I**) QPCR analysis of on effects of GSK0660 on arachidonic acid induction of *Prl8a2* mRNA level after stromal cells were treated with AA for 72h. (**J**) Western blot analysis on effects of arachidonic acid on ACTIVIN A protein level after stromal cells were treated with AA for 24 h. (**K**) QPCR analysis on effects of SB431542 (ACTIVIN A inhibitor) on arachidonic acid induction of *Prl8a2* mRNA levels. (**L**) QPCR analysis on effects of *Inhba* siRNAs on *arachidonic acid induction on Inhba* mRNA level after stromal cells were treated with arachidonic acid for 72 h. Data were presented as means ± SD from at least 3 biological replicates.

### 3.5. Blastocyst-derived TNF*α* promotes cPLA2*α* phosphorylation and arachidonic acid secretion

We just showed that arachidonic acid could initiate fibroblast activation and induce decidualization. When we examined the concentration of arachidonic acid in the uterine luminal fluid, the luminal concentration of arachidonic acid at day 4 midnight was significantly higher than that in the morning of days 3 and 4 of pregnancy, indicating AA secretion from the uterus just after embryo implantation (Fig. 5A).

**Fig. 5.**
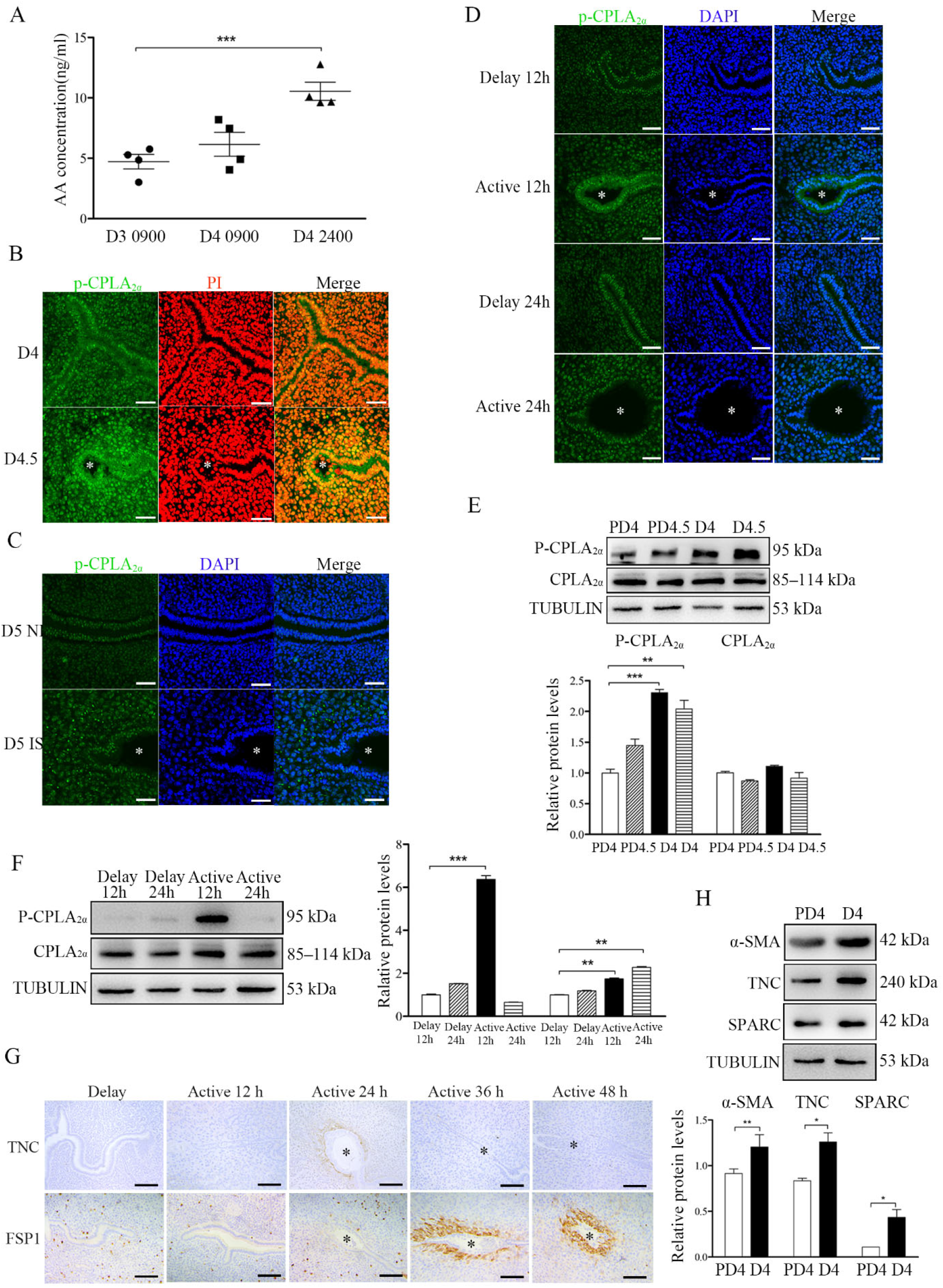
The involvement of blastocysts in fibroblast activation during early pregnancy. (**A**) Arachidonic acid concentration in uterine luminal fluid flushed on day 3 morning (10:00; n = 20 mice), day 4 morning (10:00; n=20 mice), and day 4 midnight (24:00; n = 20 mice) of pregnancy. (**B**) p-cPLA2α immunofluorescence in mouse uteri on day 4 morning (10:00; n = 6) and day 4 evening (24:00; n =6. * Embryo. Scale bar =50 μm. (**C**) p-cPLA2α immunofluorescence in mouse uteri at implantation sites and inter-implantation sites on day 5 morning (10:00; n = 6 mice) of pregnancy. * Embryo. NI, inter-implantation site; IS, implantation site. Scale bar =50 μm. (**D**) p-cPLA_2α_ immunofluorescence of in mouse uteri 12 and 24 h after delayed implantation was activated by estrogen treatment, respectively. * Embryo. Scale bar =50 μm. (**E**) Western blot analysis of cPLA_2α_ and p-cPLA_2α_ protein levels in mouse uteri on day 4 and day 4 midnight of pregnancy, and day 4 and day 4 midnight of pseudopregnancy, respectively. **(F)** Western blot analysis of cPLA2α and p-cPLA2α protein levels in mouse uteri 12 and 24 h after delayed implantation was activated by estrogen treatment. **(G)** Immunostaining of TNC and FSP1 in mouse uteri 12, 24, 36 and 48 h after delayed implantation was activated by estrogen treatment. * Embryo. **(H)** Western blot analysis α-SMA, TNC, and SPARC protein levels in mouse uteri on day 4 of pregnancy and day4 of pseudopregnancy. PD4, D4 of pseudopregnancy.

Immunofluorescence also showed that the p-cPLA2α level in luminal epithelium in day 4 midnight was obviously stronger than that on day 4 (Fig. 5B). Compare with inter-implantation sites, p-cPLA2α immunofluorescence at implantation site on D5 was also stronger (Fig. 5C). Compared with delayed implantation, p-cPLA_2α_ immunofluorescence in the luminal epithelium was stronger 12 h after estrogen activation (Fig. 5D). Western blot also confirmed that p-cPLA_2α_ levels on day 4 and day 4 midnight of pregnancy was significantly higher than day 4 and day 4 midnight of pseudopregnancy (Fig. 5E-source data-1). Compared with delayed implantation, p-cPLA2α protein levels were also higher 12 h after estrogen activation (Fig. 5F-source data-2). These results suggested that embryos should be involved in cPLA2α phosphorylation.

Furthermore, we examined the markers of fibroblast activation. Compared with delayed implantation, the immunostaining levels of both TNC and FSP1 at implantation sites after estrogen activation were stronger (Fig. 5G). Western blot also indicated that the protein levels of the markers of fibroblast activation (αSMA, TNC and SPARC) on day 4 of pregnancy were stronger than that on day 4 of pseudopregnancy (Fig. 5H-source data-3). This results strongly suggest that embryos were tightly involved in fibroblast activation.

S100A9, HB-EGF and TNFα are previously shown to be secreted by blastocysts [50–52]. When S100A9-soaked blue beads were transferred into day 4 pseudopregnant uterine lumen, FSP1 had no obvious change, but TNC immunostaining was increased slightly (Fig. 6A). When HB-EGF-soaked beads were transferred, FSP1 immunostaining was slightly increased, but TNC immunostaining was increased obviously (Fig. 6B). However, TNFα-soaked beads obviously stimulated the immunostaining levels of both FSP1 and TNC (Fig. 6C).

**Fig. 6.**
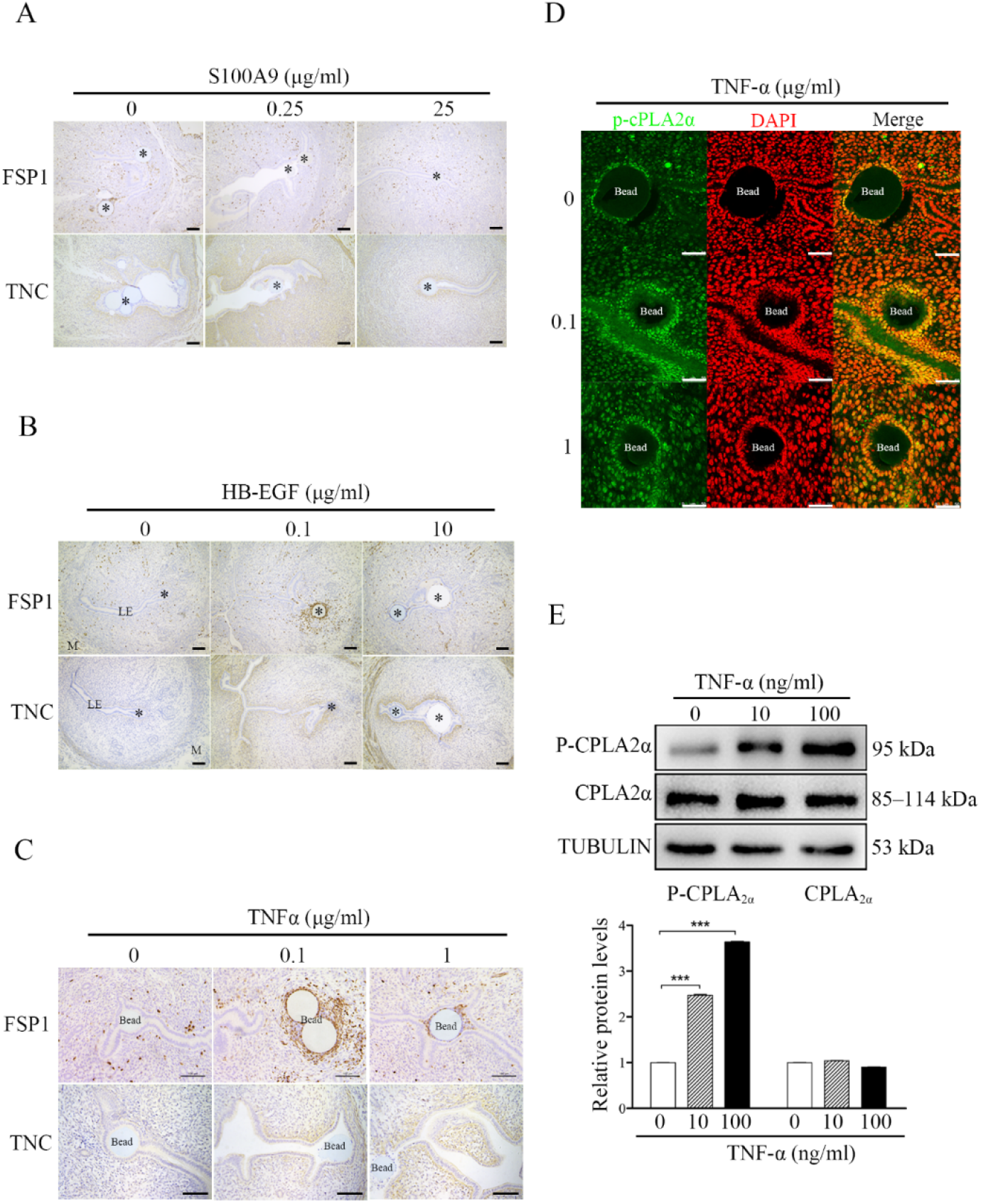
Effects of TNF*α* on cPLA_2*α*_ phosphorylation and arachidonic acid secretion. (**A**) Immunostaining of TNC and FSP1 in mouse uteri after S100A9-soaked blue beads were transferred into uterine lumen of day 4 pseudopregnant mice for 24 h. (**B**) Immunostaining of TNC and FSP1 in mouse uteri after HB-EGF-soaked blue beads were transferred into uterine lumen of day 4 pseudopregnant mice for 24 h. (**C**) Immunostaining of TNC and FSP1 in mouse uteri after TNFα-soaked blue beads were transferred into uterine lumen of day 4 pseudopregnant mice for 24 h.* Bead; LE, luminal epithelium; M, muscular layer; St, stroma. Scale bar =100 μm. (**D**) p-cPLA_2α_ immunofluorescence in mouse uteri after 0.1 and 1 μg/ml TNFα-soaked blue beads were transferred into day 4 pseudopregnant uterine lumen. Scale bar =50 μm. (**E**) Western blot analysis of cPLA_2α_ and p-cPLA_2α_ protein levels after cultured epithelial cells were treated with TNFα for 3 h.

After TNFα-soaked beads were transferred into day 4 pseudopregnant uterine lumen, p-cPLA_2α_ immunofluorescence at luminal epithelium was obviously increased (Fig. 6D). When the epithelial cells isolated day 4 pseudopregnant uterus were treated with TNFα, Western blot showed that the protein level of p-cPLA_2α_ was significantly upregulated (Fig. 6E-source data-1). These results indicated that embryo-derived TNFα was able to promote the phosphorylation of cPLA_2α_ in luminal epithelium for liberating arachidonic acid into uterine stroma.

### 3.6. Effects of fibroblast activation on TNF*α*-AA-PGI2 pathway under human in vitro decidualization

After we showed that fibroblast activation was involved in mouse decidualization, we wondered whether fibroblast activation participated in human decidualization. Compared with control group, the protein levels of α-SMA, TNC and SPARC were significantly increased under human in vitro decidualization (Fig. 7A-source data-1). Previous studied also indicated that human blastocysts could synthesize and secret TNFα [50; 53]. When human uterine epithelial ISHIKAWA cells were co-cultured with human 4003 stromal cells, TNFα treatment significantly increased the protein level of p-cPLA_2α_ in epithelial ISHIKAWA cells (Fig. 7B-source data-2), and the protein levels of TNC, αSMA, SPARC and ACTIVIN A in 4003 stromal cells (Fig. 7C-source data-3). When human stromal cells were treated with arachidonic acid, the protein levels of TNC, αSMA and SPARC were obviously stimulated (Fig. 7D-source data-4). Meanwhile, treatment with arachidonic acid also significantly upregulated the protein levels of COX-2, PGIS and PPARδ, but had no obvious effects on PGES protein level (Fig. 7E-source data-5). After 4003 stromal cells were treated with either PGI_2_ analogs ILOPROST or PPAR-δ agonists GW501516, the protein levels of TNC, αSMA and SPARC were significantly up-regulated compared with control group (Fig. 7F-source data-6, G-source data-7). These results suggested that arachidonic acid should promote fibroblast activation through PGI2– PPAR-δ pathway during human decidualization. Treatment with arachidonic acid also significantly stimulated the mRNA expression of *INHBA* (encoded for human ACTIVIN A) (Fig. 7H), which was significantly abrogated by COX-2 inhibitor NS398 (Fig. 7I). Overall, these results indicated that fibroblast activation was also involved in human decidualization in a similar mechanism as in mice.

**Fig. 7.**
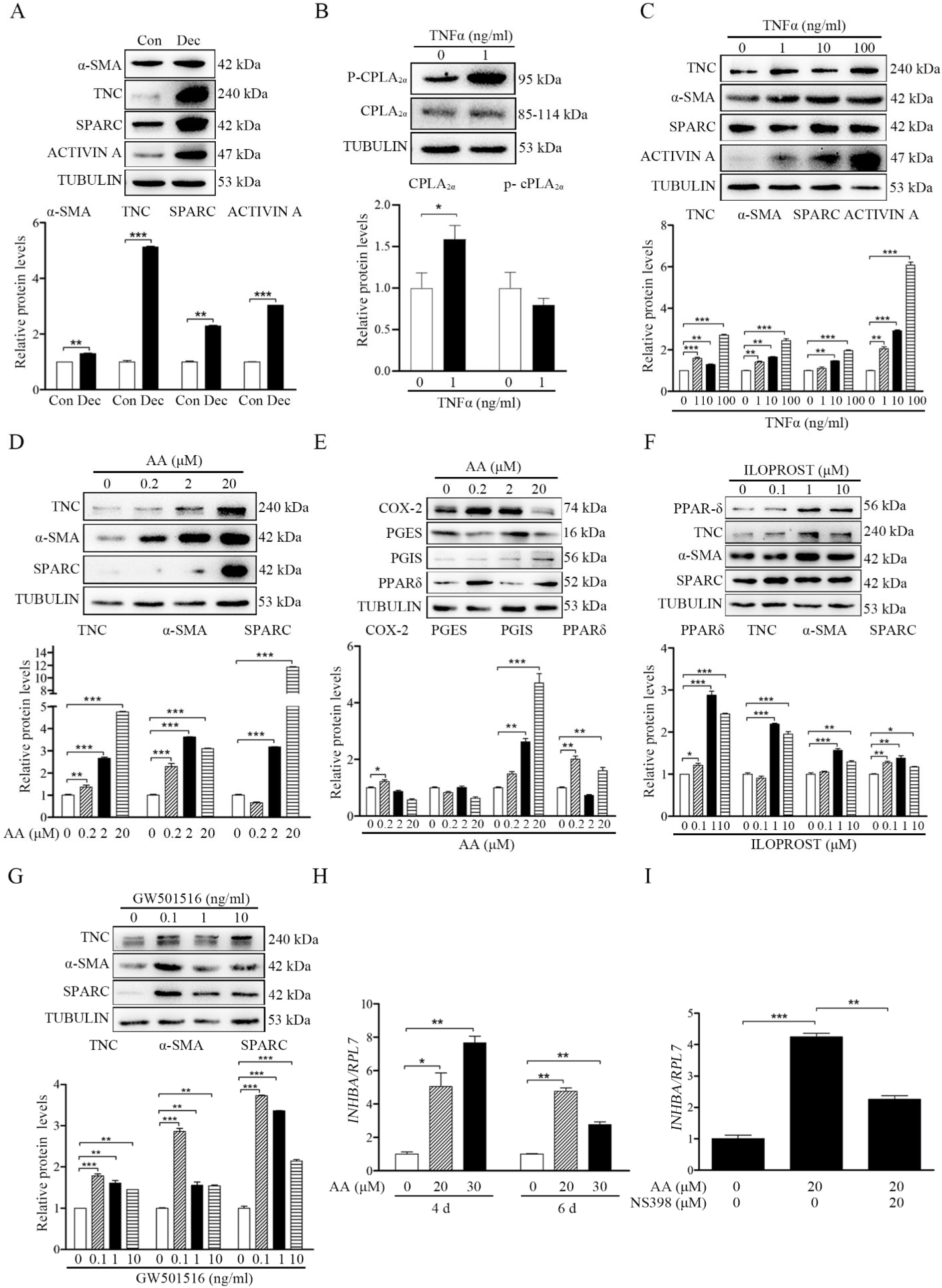
TNFα regulation of fibroblast activation through AA-PGI-ACTIVIN A pathway in human endometrium. (**A**) Western blot analysis of α-SMA, TNC, SPARC and ACTIVIN A protein levels after human stromal cells were induced for decidualization for 24 h. (**B**) Western blot analysis of cPLA2α and p-cPLA2α protein levels after human ISHIKAWA cells were treated with TNFα for 3 h. (**C**)Western blot analysis of TNC, α-SMA, SPARC and ACTIVIN A protein levels in stromal 4003 cells after the co-culture of ISHIKAWA cells and stromal cells were treated with TNFα for 3 h. (**D**) Western blot analysis of TNC, α-SMA and SPARC protein levels after stromal 4003 cells were treated with AA for 6 h. (**E**) Western blot analysis of COX-2, PGES, PGIS, and PPARδ protein levels after stromal 4003 cells were treated with AA for 3 h. (**F**) Western blot analysis of PPARδ, TNC, α-SMA and SPARC protein levels after stromal 4003 cells were treated with ILOPROST for 12 h. (**G**) Western blot analysis of TNC, α-SMA and SPARC protein levels after stromal cells 4003 cells were treated with GW501516 for 6 h. (**H**) QPCR analysis of *INHBA* mRNA levels after stromal 4003 cells were treated with AA. (**I**) QPCR analysis on effects of NS398 on arachidonic acid stimulation of *INHBA* mRNA levels after stromal 4003 cells were treated with AA.

### 3.7. Arachidonic acid-PGI2 pathway is essential to human decidualization

Insulin growth factor binding protein 1(*IGFBP1*) and prolactin (*PRL*) are recognized markers for human in vitro decidualization [54]. When human stromal cells were treated with arachidonic acid, both *IGFBP1* and *PRL* were significantly increased (Fig. 8A). Under human in vitro decidualization, either ILOPROST or GW501516 could significantly promote the mRNA expression of *IGFBP1* and *PRL* (Fig. 8B, C). The induction of arachidonic acid on *IGFBP1* and *PRL* mRNA expression was significantly abrogated by either COX-2 inhibitor NS398 or PPARδ antagonist GSK0660 (Fig. 8D). Under human in vitro decidualization, the mRNA levels of *IGFBP1* and *PRL* were also significantly up-regulated by ACTIVIN A treatment (Fig. 8E). Taken together, these results suggested that epithelium-derived arachidonic acid could promote fibroblast activation through COX-2-PGI_2_-PPAR-δ pathway and induce decidualization via ACTIVIN A during human decidualization, which was also consistent with the results in mice.

**Fig. 8.**
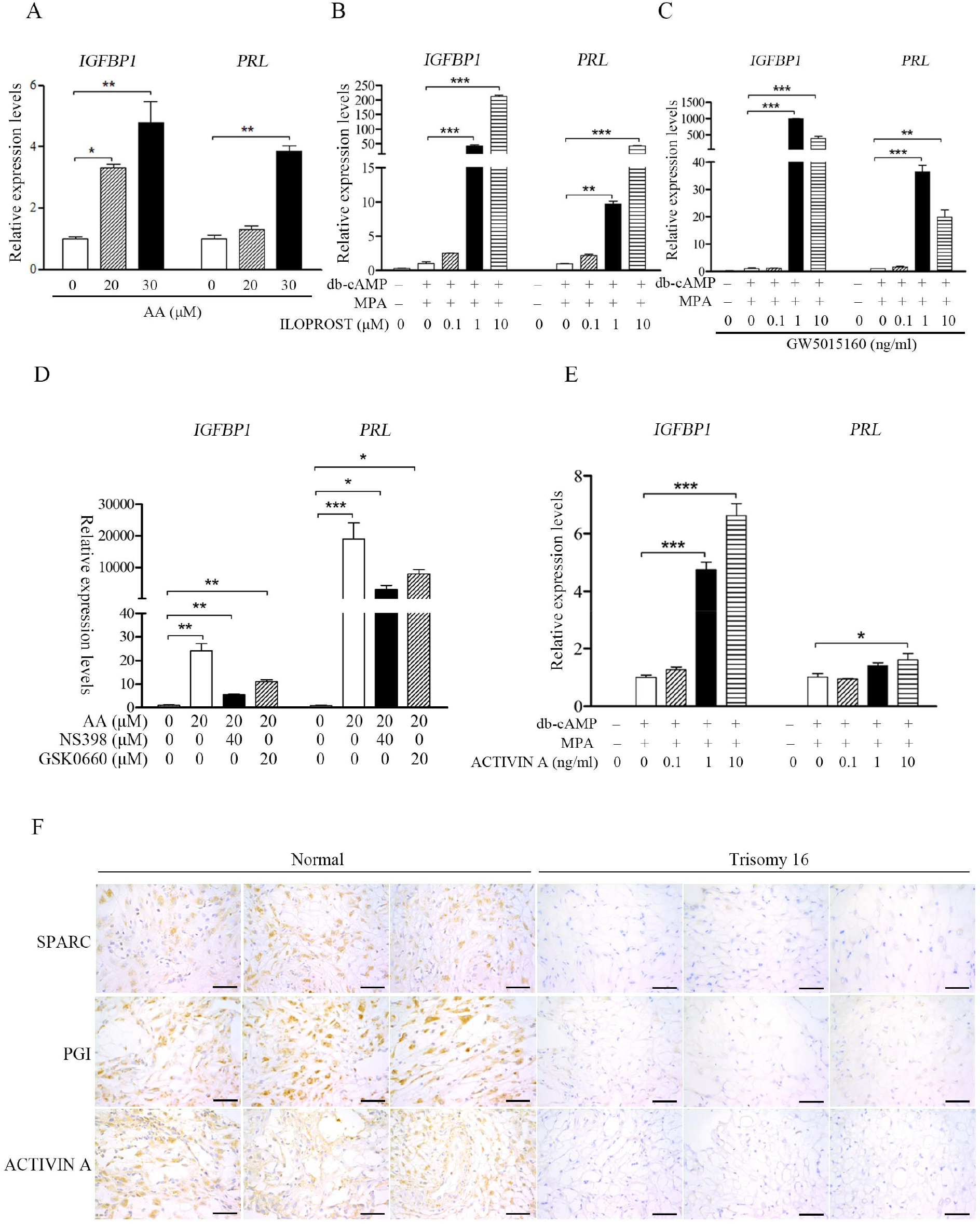
AA-PGI2-PPARδ-ACTIVIN A regulation on human decidualization. (**A**) QPCR analysis of *IGFBP1* and *PRL* mRNA levels after stromal 4003 cells were treated with AA. (**B**) QPCR analysis of *IGFBP1* and *PRL* mRNA levels after stromal 4003 cells were treated with ILOPROST for 4 days under in vitro decidualization for 4 days. (**C**) QPCR analysis of *IGFBP1* and *PRL* mRNA levels after stromal 4003 cells were treated with GW501160 for 4 days under in vitro decidualization for 4 days. (**D**) QPCR analysis on effects of NS398 or GSK0660 on AA induction of *IGFBP1* and *PRL* mRNA levels after stromal 4003 cells were treated with arachidonic acid for 4 days. (**E**) QPCR analysis of *IGFBP1* and *PRL* mRNA levels after stromal 4003 cells were treated with ACTIVIN A for 2 days under in vitro decidualization. (**F**) Immunostaining of SPARC, PGIS and ACTIVIN A in human decidual tissues from control and decidual tissues with fetal trisomy 16. Scale bar, 50 μm.

For translational study, we examined the markers of fibroblast activation in human samples. Compared with normal diploid group, the protein levels of SPARC, PGIS and ACTIVIN A in the decidual tissues with fetal trisomy 16 were obviously down-regulated (Fig. 8F), suggesting the abnormality of fibroblast activation in the decidua with fetal trisomy 16.

## 4. DISCUSSION

Our study was the first to identify that embryo-derived TNFα promotes the phosphorylation of cPLA_2α_ and the release of arachidonic acid from luminal epithelium to induce fibroblast activation and decidualization through ACTIVIN A. We also showed that the pathway underlying fibroblast activation was conserved in mice and humans.

Fibroblasts activation could be identified by many markers, including α-SMA, TNC, POSTN, NG-2, PDGF receptor-a/b, FSP1 (S100A4) and FAP. These fibroblast markers expressed alone or in combination and could be used to identify distinct subpopulations following fibroblast activation [42; 55; 56]. Fibroblast activation plays an important role under many physiological or pathological conditions, such as cancer, injury repair and fibrosis. However, the regulation and function of fibroblast activation during decidualization remain unknown. In our study, fibroblast activation was first identified during mouse and human decidualization using multiple markers. Fibroblasts are generally in a dormant and quiescent state in tissues, and are only activated when stimulated [4]. In a previous study, mPGES1-derived PGE_2_ supports the early inflammatory phase of wound healing and may stimulate subsequent fibroblast activation early after damage [57]. Although PGE_2_ has been shown to be essential for mouse decidualization [48; 58], PGE_2_ was ineffective in activating fibroblast activation. However, our study indicated that arachidonic acid from luminal epithelium was able to induce fibroblast activation via PGI2-PPARδ pathway. Previous studies indicated that arachidonic acid could promote mouse decidualization [59; 60]. COX2-derived PGI2 has been identified to be essential for mouse decidualization via PPARδ [35]. In our study, arachidonic acid concentration in uterine lumen was significantly increased during embryo attachment. Arachidonic acid, PGI2 or PPARδ agonist was able to induce fibroblast activation.

Activated fibroblasts play key roles in the injury response, tumorigenesis, fibrosis, and inflammation through secreting different factors in different physiological or pathological processes [61]. Myofibroblast marker α-SMA has been shown to be one of the important markers of early decidualization in primates[20], and sterile inflammatory secretion of products such as ATP and uric acid after injury has been shown to stimulate fibroblast activation [28; 29] and uterine decidualization [26; 27], so we speculate that fibroblast activation is strongly correlated with decidualization. Because TNC, SPARC, FSP1 and ACTIVIN A were all identified during fibroblast activation, we examined the role of each of these markers during mouse decidualization. We found that only ACTIVIN A was able to induce mouse in vitro decidualization. The function of ACTIVIN A during human decidualization was also confirmed in our study. ACTIVIN A has been shown to be important for human decidualization [62]. ACTIVIN A, its functional receptors, and binding proteins, are abundant in human endometrium [63]. Our study indicated that AA-PGI_2-_PPAR-δ axis stimulated fibroblast activation and induced decidualization through ACTIVIN A.

The adequate molecular interaction between the endometrium and the blastocyst is critical for successful implantation and decidualization [64; 65]. In our study, there was a big difference of both markers of fibroblast activation and p-cPLA2α between pregnancy and pseudopregnancy, and between delayed and activated implantation, strongly suggesting the involvement of embryos in these processes. Although S100A9, HB-EGF and TNFα are previously shown to be secreted by blastocysts [50–52], TNFα was the only one to stimulate both FSP1 and TNC, and to phosphorylate cPLA2α in our study. Cytosolic phospholipase A2α (cPLA2α, encoded by *Pla2g4a*) is a major provider of arachidonic acid (AA). *Pla2g4a*– /–mice results in deferred implantation and deranged gestational development [33]. We also showed that arachidonic acid concentration in luminal fluid was significantly increased in day 4 evening when the embryos just implanted. TNFα is present in the reactivated blastocyst and human blastocyst [50; 53], and may play a critical role during embryo implantation [66]. In our study, we confirmed that TNFα stimulated the phosphorylation of cPLA2α and arachidonic acid release from luminal epithelium. A proper interaction between embryos and decidualization is critical for successful pregnancy. It is shown that impaired decidualization from recurrent pregnancy loss is unable to distinguish the quality of implanting blastocysts [67].

Additionally, the quality of blastocysts is essential for uNK cells to kill senescent decidual cells [68]. There is a high incidence of chromosome aneuploidy in human gametes and embryos, which is a major cause of implantation failure and miscarriage [69]. In the trisomy 18 pregnancies, the fetal and maternal hCG values were significantly lower than in controls. However, in Turner syndrome pregnancies, both fetal and maternal values were significantly higher than in controls [70]. Indeed, we found that fibroblast activation was impaired in the decidual tissues with fetal 16 trisomy. It is interesting to note that our experimental results show that the regulatory mechanism and function of fibroblast activation are conserved in humans and mice.

Activated fibroblasts play an important role in many physiological and pathologic processes. Excessive fibroblast activation can lead to fibrosis [46; 71; 72]. Fibroblast activation may be present and balanced during normal pregnancy without ultimately leading to fibrosis or other diseases. S100A4 is hypomethylated and overexpressed in grade 3 endometrioid tumors compared with benign endometrium [73]. Genetic and proteomic analysis of surgical specimens from 14 patients with uterine leiomyoma showed that TNC is significantly upregulated in patient samples [74]. Fibrosis mostly occurs in pathological conditions. Intrauterine adhesions (IUA), also known as Ashman syndrome, are caused by endometrial fibrosis as a result of injury to the uterus’s basal lining, resulting in partial or total adhesions in the uterine cavity [75]. IUA can interfere with the embryo’s implantation and development, resulting in decreased or even full loss of intrauterine volume, female infertility, and recurrent miscarriages [76]. The thin endometrial model exhibits a higher degree of fibrosis than normal controls, which is thought to be a crucial component in embryo implantation failure [77]. The key question is why fibrosis doesn’t occur during normal early pregnancy? Both activins and inhibins are expressed in pregnant uterus [78]. In our study, *Fst* was significantly up-regulated under in vitro decidualization although ACTIVIN A was able to stimulate in vitro decidualization. FST is a secreted glycoprotein and can neutralize the profibrotic and proinflammatory actions of ACTIVINS. FST has a strong antifibrotic effect in various organs [79; 80]. Furthermore, FST is shown to be critical for mouse decidualization [81]. Additionally, arachidonic acid was able to induce fibroblast activation and promote decidualization in our study. A recent study showed that 11,12-epoxyeicosatrienoic acid, a metabolite of arachidonic acid can alleviate pulmonary fibrosis [82]. It is possible that a physiological level of fibroblast activation is beneficial for decidualization and the long-lasting fibroblast activation could be balanced by certain molecules, like FST or arachidonic acid metabolite.

## 5. CONCLUSION

In this study, we identified that embryos-derived TNFα was able to phosphorylate cPLA2α for releasing arachidonic acid from luminal epithelium. Arachidonic acid could physiologically induce fibroblast activation and promote decidualization via PGI2-PPARδ-ACTIVIN A axis. This regulatory mechanism was also conserved in mice and humans. Overall, this study should shed a light on the novel mechanism underlying decidualization.

## 6. ABBREVIATIONS

AA: Arachidonic acid
CAFs: Cancer-associated fibroblasts
DAMPs: Damage associated molecular patterns
ECM: Extracellular matrix
FA: Fibroblast activation
IUA: Intrauterine adhesions
POSTN: Periostin
TNC: Tenascin C

## 7. DECLARATIONS

### Ethics approval and consent to participate

This study was approved by The Ethics Committee of Zhejiang University School of Medicine’s Obstetrics and Gynecology Hospital and the Human Research Committee of Nanjing Drum Tower Hospital, respectively. All animal protocols were approved by the Animal Care and Use Committee of South China Agricultural University.

### Consent for publication

Not applicable.

### Availability of data and materials

All data are available in the main text.

### Competing interests

The authors declare that they have no competing interests

### Funding

This study was supported by the National Key Research and Development Program of China (2018YFC1004400) and National Natural Science Foundation of China (31871511 and 32171114)

### Author contributions

Design experiment: STC, ZMY

Experiments performed: STC, WWS, YQL, WY, LY

Data analysis: STC, ZSY, MYL, ZMY

Provide clinical samples: AXL, YLH

Writing – manuscript: STC, ZMY

Writing – review & editing: STC, ZMY, YLH

All authors read and approved the final manuscript.

## Acknowledgments

Not applicable.

## Legends for uncropped Western blot images

Fig. 1B-source data-1. Western blot analysis of α-SMA, SPARC, TNC protein level under in vitro decidualization (EP) for 24 h.

Fig. 2A-source data-1. Western blot analysis on the effects of TNC on decidualization markers (BMP2, WNT4, E2F8 and CYCLIN D3) after stromal cells were treatment with TNC for 72 h.

Fig. 2B-source data-2. Western blot analysis of the effects of FSP1 on decidualization markers after stromal cells were treated with FSP1 for 72 h.

Fig. 2C-source data-3. Western blot analysis on the effects of Sparc overexpression on decidualization markers after overexpression of Sparc gene in cultured stromal cells.

Fig. 2D-source data-4. Western blot analysis on ACTIVIN A protein levels in mouse uteri on day 4 0900 and day 4 2400 of pregnancy and pseudopregnancy, respectively.

Fig. 2E-source data-5. Western blot analysis on the effects of ACTIVIN A on decidualization markers after stromal cells were treated with ACTIVIN A for 72 h.

Fig. 2F-source data-6. Western blot analysis on the effects of ACTIVIN A on decidualization markers after stromal cells were treated with ACTIVIN A for 48 h under in vitro decidualization.

Fig. 3A-source data-1. The effects of PGE2 on markers of fibroblast activation.

Fig. 3B-source data-2. The effects of ILOPROST, PGI2 analog, on markers of fibroblast activation after stromal cells was treated with PGI2 for 12 h.

Fig. 3C-source data-3. The effects of GW501516, PPARδ agonist, on markers of fibroblast activation.

Fig. 3D-source data-4. The effects of ILOPROST on decidualization markers.

Fig. 3E-source data-5. The effects of GW501516 on decidualization markers.

Fig. 3F-source data-6. The effects of ILOPROST on ACTIVIN A protein levels after stromal cells were treated with ILOPROST for 24 h.

Fig. 4A-source data-1. Western blot analysis on effects of arachidonic acid on markers of fibroblast activation after stromal cells were treated with arachidonic acid for 6 h.

Fig. 4B-source data-2. Western blot analysis on effects of arachidonic acid on decidualization markers after stromal cells were treated with arachidonic acid for 48 h.

Fig. 4C-source data-3. Western blot analysis on effects of arachidonic acid on decidualization markers after stromal cells were treated with arachidonic acid for 48 h under in vitro decidualization. EP, 17β-estradiol + progesterone.

Fig. 4E-source data-4. Western blot analysis on effects of arachidonic acid on COX2, PGES, PGIS and PPARδ protein levels after stromal cells were treated with arachidonic acid for 6 h.

Fig. 4F-source data-5. Western blot analysis on effects of NS398 (COX-2 inhibitor) on arachidonic acid induction of COX2, PGES, PGIS and PPARδ protein levels after stromal cells were treated with arachidonic acid for 48 h in the absence or presence of NS398.

Fig. 4J-source data-6. Western blot analysis on effects of arachidonic acid on ACTIVIN A protein level after stromal cells were treated with AA for 24 h.

Fig. 5E-source data-1. Western blot analysis of cPLA2α and p-cPLA2α protein levels in mouse uteri on day 4 and day 4 midnight of pregnancy, and day 4 and day 4 midnight of pseudopregnancy, respectively.

Fig. 5F-source data-2. Western blot analysis of cPLA2α and p-cPLA2α protein levels in mouse uteri 12 and 24 h after delayed implantation was activated by estrogen treatment.

Fig. 5H-source data-3. Western blot analysis α-SMA, TNC, and SPARC protein levels in mouse uteri on day 4 of pregnancy and day4 of pseudopregnancy.

Fig. 6E-source data-1. Western blot analysis of cPLA2α and p-cPLA2α protein levels after cultured epithelial cells were treated with TNFα for 3 h.

Fig. 7A-source data-1. Western blot analysis of α-SMA, TNC, SPARC and ACTIVIN A protein levels after human stromal cells were induced for decidualization for 24 h.

Fig. 7B-source data-2. Western blot analysis of cPLA2α and p-cPLA2α protein levels after human ISHIKAWA cells were treated with TNFα for 3 h.

Fig. 7C-source data-3. Western blot analysis of TNC, α-SMA, SPARC and ACTIVIN A protein levels in stromal 4003 cells after the co-culture of ISHIKAWA cells and stromal cells were treated with TNFα for 3 h.

Fig. 7D-source data-4. Western blot analysis of TNC, α-SMA and SPARC protein levels after stromal 4003 cells were treated with AA for 6 h.

Fig. 7E-source data-5. Western blot analysis of COX-2, PGES, PGIS, and PPARδ protein levels after stromal 4003 cells were treated with AA for 3 h.

Fig. 7F-source data-6. Western blot analysis of PPARδ, TNC, α-SMA and SPARC protein levels after stromal 4003 cells were treated with ILOPROST for 12 h.

Fig. 7G-source data-7. Western blot analysis of TNC, α-SMA and SPARC protein levels after stromal cells 4003 cells were treated with GW501516 for 6 h.

## Notes

### Competing Interest Statement

The authors have declared no competing interest.

## REFERENCES

[1] Yoshida, G.J., 2020. Regulation of heterogeneous cancer-associated fibroblasts: the molecular pathology of activated signaling pathways. J Exp Clin Cancer Res 39(1):112.

[2] Tracy, L.E., Minasian, R.A., Caterson, E.J., 2016. Extracellular Matrix and Dermal Fibroblast Function in the Healing Wound. Adv Wound Care (New Rochelle) 5(3):119–136.

[3] Enzerink, A., Vaheri, A., 2011. Fibroblast activation in vascular inflammation. J Thromb Haemost 9(4):619–626.

[4] Pakshir, P., Noskovicova, N., Lodyga, M., Son, D.O., Schuster, R., Goodwin, A., et al., 2020. The myofibroblast at a glance. J Cell Sci 133(13).

[5] Tomasek, J.J., Gabbiani, G., Hinz, B., Chaponnier, C., Brown, R.A., 2002. Myofibroblasts and mechano-regulation of connective tissue remodelling. Nature Reviews Molecular Cell Biology 3(5):349–363.

[6] Angelini, A., Trial, J., Ortiz-Urbina, J., Cieslik, K.A., 2020. Mechanosensing dysregulation in the fibroblast: A hallmark of the aging heart. Ageing Res Rev 63:101150.

[7] Nurmik, M., Ullmann, P., Rodriguez, F., Haan, S., Letellier, E., 2020. In search of definitions: Cancer-associated fibroblasts and their markers. Int J Cancer 146(4):895–905.

[8] Shimura, T., 2021. Roles of Fibroblasts in Microenvironment Formation Associated with Radiation-Induced Cancer. Adv Exp Med Biol 1329:239–251.

[9] Kuzet, S.E., Gaggioli, C., 2016. Fibroblast activation in cancer: when seed fertilizes soil. Cell Tissue Res 365(3):607–619.

[10] Martin, R.D., 2007. The evolution of human reproduction: a primatological perspective. Am J Phys Anthropol Suppl 45:59–84.

[11] Salamonsen, L.A., Hutchison, J.C., Gargett, C.E., 2021. Cyclical endometrial repair and regeneration. Development 148(17).

[12] Schuster, R., Rockel, J.S., Kapoor, M., Hinz, B., 2021. The inflammatory speech of fibroblasts. Immunol Rev 302(1):126–146.

[13] Wang, W., Vilella, F., Alama, P., Moreno, I., Mignardi, M., Isakova, A., et al., 2020. Single-cell transcriptomic atlas of the human endometrium during the menstrual cycle. Nat Med 26(10):1644–1653.

[14] Lv, H., Zhao, G., Jiang, P., Wang, H., Wang, Z., Yao, S., et al., 2022. Deciphering the endometrial niche of human thin endometrium at single-cell resolution. Proc Natl Acad Sci U S A 119(8).

[15] Wang, H., Dey, S.K., 2006. Roadmap to embryo implantation: clues from mouse models. Nat Rev Genet 7(3):185–199.

[16] Li, Y., Sun, X., Dey, S.K., 2015. Entosis allows timely elimination of the luminal epithelial barrier for embryo implantation. Cell Rep 11(3):358–365.

[17] Gellersen, B., Brosens, J.J., 2014. Cyclic decidualization of the human endometrium in reproductive health and failure. Endocr Rev 35(6):851–905.

[18] Tan, Y., Li, M., Cox, S., Davis, M.K., Tawfik, O., Paria, B.C., et al., 2004. HB-EGF directs stromal cell polyploidy and decidualization via cyclin D3 during implantation. Dev Biol 265(1):181–195.

[19] McConaha, M.E., Eckstrum, K., An, J., Steinle, J.J., Bany, B.M., 2011. Microarray assessment of the influence of the conceptus on gene expression in the mouse uterus during decidualization. Reproduction 141(4):511–527.

[20] Kim, J.J., Jaffe, R.C., Fazleabas, A.T., 1999. Blastocyst invasion and the stromal response in primates. Hum Reprod 14 Suppl 2:45–55.

[21] Venuto, L., Lindsay, L.A., Murphy, C.R., 2008. Moesin is involved in the cytoskeletal remodelling of rat decidual cells. Acta Histochem 110(6):491–496.

[22] Fazleabas, A.T., Donnelly, K.M., Srinivasan, S., Fortman, J.D., Miller, J.B., 1999. Modulation of the baboon (Papio anubis) uterine endometrium by chorionic gonadotrophin during the period of uterine receptivity. Proc Natl Acad Sci U S A 96(5):2543–2548.

[23] Fujigaki, Y., Muranaka, Y., Sun, D., Goto, T., Zhou, H., Sakakima, M., et al., 2005. Transient myofibroblast differentiation of interstitial fibroblastic cells relevant to tubular dilatation in uranyl acetate-induced acute renal failure in rats. Virchows Arch 446(2):164–176.

[24] Chen, G.Y., Nuñez, G., 2010. Sterile inflammation: sensing and reacting to damage. Nat Rev Immunol 10(12):826–837.

[25] Venugopal, H., Hanna, A., Humeres, C., Frangogiannis, N.G., 2022. Properties and Functions of Fibroblasts and Myofibroblasts in Myocardial Infarction. Cells 11(9).

[26] Gu, X.W., Chen, Z.C., Yang, Z.S., Yang, Y., Yan, Y.P., Liu, Y.F., et al., 2020. Blastocyst-induced ATP release from luminal epithelial cells initiates decidualization through the P2Y2 receptor in mice. Sci Signal 13(646).

[27] Zhu, Y.Y., Wu, Y., Chen, S.T., Kang, J.W., Pan, J.M., Liu, X.Z., et al., 2021. In situ Synthesized Monosodium Urate Crystal Enhances Endometrium Decidualization via Sterile Inflammation During Pregnancy. Front Cell Dev Biol 9:702590.

[28] Dolmatova, E., Spagnol, G., Boassa, D., Baum, J.R., Keith, K., Ambrosi, C., et al., 2012. Cardiomyocyte ATP release through pannexin 1 aids in early fibroblast activation. Am J Physiol Heart Circ Physiol 303(10):H1208–1218.

[29] Bao, J., Shi, Y., Tao, M., Liu, N., Zhuang, S., Yuan, W., 2018. Pharmacological inhibition of autophagy by 3-MA attenuates hyperuricemic nephropathy. Clin Sci (Lond) 132(21):2299–2322.

[30] Baiocchini, A., Montaldo, C., Conigliaro, A., Grimaldi, A., Correani, V., Mura, F., et al., 2016. Extracellular Matrix Molecular Remodeling in Human Liver Fibrosis Evolution. PLoS One 11(3):e0151736.

[31] Yu, D., Wong, Y.M., Cheong, Y., Xia, E., Li, T.C., 2008. Asherman syndrome--one century later. Fertil Steril 89(4):759–779.

[32] Bazoobandi, S., Tanideh, N., Rahmanifar, F., Zare, S., Koohi-Hosseinabadi, O., Razeghian-Jahromi, I., et al., 2020. Preventive Effects of Intrauterine Injection of Bone Marrow-Derived Mesenchymal Stromal Cell-Conditioned Media on Uterine Fibrosis Immediately after Endometrial Curettage in Rabbit. Stem Cells Int 2020:8849537.

[33] Song, H., Lim, H., Paria, B.C., Matsumoto, H., Swift, L.L., Morrow, J., et al., 2002. Cytosolic phospholipase A2alpha is crucial [correction of A2alpha deficiency is crucial] for ‘on-time’ embryo implantation that directs subsequent development. Development 129(12):2879–2889.

[34] Lim, H., Dey, S.K., 1997. Prostaglandin E2 receptor subtype EP2 gene expression in the mouse uterus coincides with differentiation of the luminal epithelium for implantation. Endocrinology 138(11):4599–4606.

[35] Lim, H., Gupta, R.A., Ma, W.G., Paria, B.C., Moller, D.E., Morrow, J.D., et al., 1999. Cyclo-oxygenase-2-derived prostacyclin mediates embryo implantation in the mouse via PPARdelta. Genes Dev 13(12):1561–1574.

[36] Wang, H., Xie, H., Sun, X., Tranguch, S., Zhang, H., Jia, X., et al., 2007. Stage-specific integration of maternal and embryonic peroxisome proliferator-activated receptor delta signaling is critical to pregnancy success. J Biol Chem 282(52):37770–37782.

[37] Li, S.Y., Song, Z., Yan, Y.P., Li, B., Song, M.J., Liu, Y.F., et al., 2020. Aldosterone from endometrial glands is benefit for human decidualization. Cell Death Dis 11(8):679.

[38] Zheng, H.T., Zhang, H.Y., Chen, S.T., Li, M.Y., Fu, T., Yang, Z.M., 2020. The detrimental effects of stress-induced glucocorticoid exposure on mouse uterine receptivity and decidualization. FASEB J 34(11):14200–14216.

[39] Fu, T., Zheng, H.T., Zhang, H.Y., Chen, Z.C., Li, B., Yang, Z.M., 2019. Oncostatin M expression in the mouse uterus during early pregnancy promotes embryo implantation and decidualization. FEBS Lett 593(15):2040–2050.

[40] Nallasamy, S., Li, Q., Bagchi, M.K., Bagchi, I.C., 2012. Msx homeobox genes critically regulate embryo implantation by controlling paracrine signaling between uterine stroma and epithelium. PLoS Genet 8(2):e1002500.

[41] Liang, X.H., Deng, W.B., Li, M., Zhao, Z.A., Wang, T.S., Feng, X.H., et al., 2014. Egr1 protein acts downstream of estrogen-leukemia inhibitory factor (LIF)-STAT3 pathway and plays a role during implantation through targeting Wnt4. J Biol Chem 289(34):23534–23545.

[42] Prime, S.S., Cirillo, N., Hassona, Y., Lambert, D.W., Paterson, I.C., Mellone, M., et al., 2017. Fibroblast activation and senescence in oral cancer. J Oral Pathol Med 46(2):82–88.

[43] Lee, K.Y., Jeong, J.W., Wang, J., Ma, L., Martin, J.F., Tsai, S.Y., et al., 2007. Bmp2 is critical for the murine uterine decidual response. Mol Cell Biol 27(15):5468–5478.

[44] Li, Q., Kannan, A., Wang, W., Demayo, F.J., Taylor, R.N., Bagchi, M.K., et al., 2007. Bone morphogenetic protein 2 functions via a conserved signaling pathway involving Wnt4 to regulate uterine decidualization in the mouse and the human. J Biol Chem 282(43):31725–31732.

[45] Qi, Q.R., Zhao, X.Y., Zuo, R.J., Wang, T.S., Gu, X.W., Liu, J.L., et al., 2015. Involvement of atypical transcription factor E2F8 in the polyploidization during mouse and human decidualization. Cell Cycle 14(12):1842–1858.

[46] Truffi, M., Sorrentino, L., Corsi, F., 2020. Fibroblasts in the Tumor Microenvironment. Adv Exp Med Biol 1234:15–29.

[47] Loomans, H.A., Andl, C.D., 2014. Intertwining of Activin A and TGFβ Signaling: Dual Roles in Cancer Progression and Cancer Cell Invasion. Cancers (Basel) 7(1):70–91.

[48] Holmes, P.V., Gordashko, B.J., 1980. Evidence of prostaglandin involvement in blastocyst implantation. J Embryol Exp Morphol 55:109–122.

[49] Simmons, D.L., Botting, R.M., Hla, T., 2004. Cyclooxygenase isozymes: the biology of prostaglandin synthesis and inhibition. Pharmacol Rev 56(3):387–437.

[50] He, B., Zhang, H., Wang, J., Liu, M., Sun, Y., Guo, C., et al., 2019. Blastocyst activation engenders transcriptome reprogram affecting X-chromosome reactivation and inflammatory trigger of implantation. Proc Natl Acad Sci U S A 116(33):16621–16630.

[51] Haller, M., Yin, Y., Ma, L., 2019. Development and utilization of human decidualization reporter cell line uncovers new modulators of female fertility. Proc Natl Acad Sci U S A 116(39):19541–19551.

[52] Hamatani, T., Daikoku, T., Wang, H., Matsumoto, H., Carter, M.G., Ko, M.S., et al., 2004. Global gene expression analysis identifies molecular pathways distinguishing blastocyst dormancy and activation. Proc Natl Acad Sci U S A 101(28):10326–10331.

[53] Lv, J., Shan, X., Yang, H., Wen, Y., Zhang, X., Chen, H., et al., 2022. Single Cell Proteomics Profiling Reveals That Embryo-Secreted TNF-α Plays a Critical Role During Embryo Implantation to the Endometrium. Reprod Sci 29(5):1608–1617.

[54] Saleh, L., Otti, G.R., Fiala, C., Pollheimer, J., Knöfler, M., 2011. Evaluation of human first trimester decidual and telomerase-transformed endometrial stromal cells as model systems of in vitro decidualization. Reprod Biol Endocrinol 9:155.

[55] Grinnell, F., 1994. Fibroblasts, myofibroblasts, and wound contraction. J Cell Biol 124(4):401–404.

[56] Oliver, C., Montes, M.J., Galindo, J.A., Ruiz, C., Olivares, E.G., 1999. Human decidual stromal cells express alpha-smooth muscle actin and show ultrastructural similarities with myofibroblasts. Hum Reprod 14(6):1599–1605.

[57] Stratton, R., Shiwen, X., 2010. Role of prostaglandins in fibroblast activation and fibrosis. J Cell Commun Signal 4(2):75–77.

[58] Baskar, J.F., Torchiana, D.F., Biggers, J.D., Corey, E.J., Andersen, N.H., Subramanian, N., 1981. Inhibition of hatching of mouse blastocysts in vitro by various prostaglandin antagonists. J Reprod Fertil 63(2):359–363.

[59] Tessier-Prigent, A., Willems, R., Lagarde, M., Garrone, R., Cohen, H., 1999. Arachidonic acid induces differentiation of uterine stromal to decidual cells. European Journal of Cell Biology 78(6):398–406.

[60] Handwerger, S., Barry, S., Barrett, J., Markoff, E., Zeitler, P., Cwikel, B., et al., 1981. Inhibition of the synthesis and secretion of decidual prolactin by arachidonic acid. Endocrinology 109(6):2016–2021.

[61] Avery, D., Govindaraju, P., Jacob, M., Todd, L., Monslow, J., Pure, E., 2018. Extracellular matrix directs phenotypic heterogeneity of activated fibroblasts. Matrix Biol 67:90–106.

[62] Zhao, H.J., Chang, H.M., Zhu, H., Klausen, C., Li, Y., Leung, P.C.K., 2018. Bone Morphogenetic Protein 2 Promotes Human Trophoblast Cell Invasion by Inducing Activin A Production. Endocrinology 159(7):2815–2825.

[63] He, Z.Y., Liu, H.C., Mele, C.A., Barmat, L., Veeck, L.L., Davis, O., et al., 1999. Expression of inhibin/activin subunits and their receptors and binding proteins in human preimplantation embryos. J Assist Reprod Genet 16(2):73–80.

[64] Massimiani, M., Lacconi, V., La Civita, F., Ticconi, C., Rago, R., Campagnolo, L., 2019. Molecular Signaling Regulating Endometrium-Blastocyst Crosstalk. Int J Mol Sci 21(1).

[65] Latifi, Z., Fattahi, A., Ranjbaran, A., Nejabati, H.R., Imakawa, K., 2018. Potential roles of metalloproteinases of endometrium-derived exosomes in embryo-maternal crosstalk during implantation. J Cell Physiol 233(6):4530–4545.

[66] You, Y., Stelzl, P., Joseph, D.N., Aldo, P.B., Maxwell, A.J., Dekel, N., et al., 2021. TNF-α Regulated Endometrial Stroma Secretome Promotes Trophoblast Invasion. Front Immunol 12:737401.

[67] Salker, M., Teklenburg, G., Molokhia, M., Lavery, S., Trew, G., Aojanepong, T., et al., 2010. Natural selection of human embryos: impaired decidualization of endometrium disables embryo-maternal interactions and causes recurrent pregnancy loss. PLoS One 5(4):e10287.

[68] Kong, C.S., Ordoñez, A.A., Turner, S., Tremaine, T., Muter, J., Lucas, E.S., et al., 2021. Embryo biosensing by uterine natural killer cells determines endometrial fate decisions at implantation. FASEB J 35(4):e21336.

[69] Kimelman, D., Pavone, M.E., 2021. Non-invasive prenatal testing in the context of IVF and PGT-A. Best Pract Res Clin Obstet Gynaecol 70:51–62.

[70] Abbas, A., Chard, T., Nicolaides, K., 1995. Fetal and maternal hCG concentration in aneuploid pregnancies. Br J Obstet Gynaecol 102(7):561–563.

[71] Giusti, I., Di Francesco, M., Poppa, G., Esposito, L., D’Ascenzo, S., Dolo, V., 2022. Tumor-Derived Extracellular Vesicles Activate Normal Human Fibroblasts to a Cancer-Associated Fibroblast-Like Phenotype, Sustaining a Pro-Tumorigenic Microenvironment. Front Oncol 12:839880.

[72] Yeo, S.Y., Lee, K.W., Shin, D., An, S., Cho, K.H., Kim, S.H., 2018. A positive feedback loop bi-stably activates fibroblasts. Nat Commun 9(1):3016.

[73] Xie, R., Loose, D.S., Shipley, G.L., Xie, S., Bassett, R.L., Jr., Broaddus, R.R., 2007. Hypomethylation-induced expression of S100A4 in endometrial carcinoma. Mod Pathol 20(10):1045–1054.

[74] Jamaluddin, M.F.B., Nagendra, P.B., Nahar, P., Oldmeadow, C., Tanwar, P.S., 2019. Proteomic Analysis Identifies Tenascin-C Expression Is Upregulated in Uterine Fibroids. Reprod Sci 26(4):476–486.

[75] Healy, M.W., Schexnayder, B., Connell, M.T., Terry, N., DeCherney, A.H., Csokmay, J.M., et al., 2016. Intrauterine adhesion prevention after hysteroscopy: a systematic review and meta-analysis. Am J Obstet Gynecol 215(3):267–275.e267.

[76] Leung, R.K., Lin, Y., Liu, Y., 2021. Recent Advances in Understandings Towards Pathogenesis and Treatment for Intrauterine Adhesion and Disruptive Insights from Single-Cell Analysis. Reprod Sci 28(7):1812–1826.

[77] Zhang, L., Li, Y., Dong, Y.C., Guan, C.Y., Tian, S., Lv, X.D., et al., 2022. Transplantation of umbilical cord-derived mesenchymal stem cells promotes the recovery of thin endometrium in rats. Sci Rep 12(1):412.

[78] Ni, N., Li, Q., 2017. TGFβ superfamily signaling and uterine decidualization. Reprod Biol Endocrinol 15(1):84.

[79] Aoki, F., Kurabayashi, M., Hasegawa, Y., Kojima, I., 2005. Attenuation of bleomycin-induced pulmonary fibrosis by follistatin. Am J Respir Crit Care Med 172(6):713–720.

[80] Patella, S., Phillips, D.J., Tchongue, J., de Kretser, D.M., Sievert, W., 2006. Follistatin attenuates early liver fibrosis: effects on hepatic stellate cell activation and hepatocyte apoptosis. Am J Physiol Gastrointest Liver Physiol 290(1):G137–144.

[81] Fullerton, P.T., Jr., Monsivais, D., Kommagani, R., Matzuk, M.M., 2017. Follistatin is critical for mouse uterine receptivity and decidualization. Proc Natl Acad Sci U S A 114(24):E4772–e4781.

[82] Kim, H.S., Moon, S.J., Lee, S.E., Hwang, G.W., Yoo, H.J., Song, J.W., 2021. The arachidonic acid metabolite 11,12-epoxyeicosatrienoic acid alleviates pulmonary fibrosis. Exp Mol Med 53(5):864–874.

